# The GPIbα intracellular tail - role in transducing VWF- and Collagen/GPVI-mediated signaling

**DOI:** 10.1101/2020.12.16.423021

**Authors:** Adela Constantinescu-Bercu, Yuxiao Alice Wang, Kevin J Woollard, Pierre Mangin, Karen Vanhoorelbeke, James TB Crawley, Isabelle I. Salles-Crawley

## Abstract

Synergy between GPIbα and GPVI signaling machineries has been suggested previously, however its molecular mechanism remains unclear. We generated a novel *GPIbα* transgenic mouse (*GPIbα*^Δsig/Δsig^) by CRISPR-Cas9 technology to delete the last 24 residues of the GPIbα intracellular tail important for VWF-mediated signaling. *GPIbα*^Δsig/Δsig^ platelets bound VWF normally under flow but formed fewer filopodia on VWF/botrocetin, demonstrating that the deleted region does not affect ligand binding but appreciably impairs VWF-dependent signaling. Notably, while haemostasis was normal in *GPIbα*^Δsig/Δsig^ mice, *GPIbα*^Δsig/Δsig^ platelets exhibited defective responses after collagen-related-peptide stimulation and formed smaller aggregates on collagen-coated microchannels at low and high shears. Flow assays performed with plasma-free blood or in the presence of α_IIb_β_3_-or GPVI-blockers suggested reduced α_IIb_β_3_ activation contributes to the phenotype of the *GPIbα*^Δsig/Δsig^ platelets. Together, these results reveal a new role for the intracellular tail of GPIbα in transducing both VWF-GPIbα and collagen-GPVI signaling events in platelets.

**Summary statement:** GPIbα and GPVI are two key receptors on the platelet surface. Using a novel transgenic mouse (*GPIbα*^Δsig/Δsig^) that lacks the last 24 amino acids of the GPIbα intracellular tail, we demonstrate the importance of this region not only in transducing signals in response to GPIbα binding to VWF, but also for collagen-GPVI-mediated platelet responses revealing previously underappreciated receptor crosstalk between GPIbα and GPVI.

## Introduction

Platelets play crucial roles in hemostasis, as well as several other physiological systems. To fulfil their hemostatic function, platelets are recruited to sites of vessel damage by von Willebrand factor (VWF), which interacts both with exposed collagen (via its A3 domain) and platelets, through specific binding of its A1 domain to glycoprotein (GP) Ibα on the platelet surface. Shear-dependent VWF-mediated capture of platelets is transient in nature, allowing platelets to roll on VWF surfaces due to the fast association and dissociation rates of the VWF A1-GPIbα interaction.(Li, 2018) Interaction of platelets with additional ligands (e.g. α_IIb_β_3_-fibrinogen, collagen-GPVI, collagen-α_2_β_1_) as well as changes in platelet phenotype are required to stabilize platelet binding to the damaged vessel wall. Although the VWF-GPIbα interaction primarily facilitates platelet recruitment under high shear conditions, it also modifies platelet phenotype through induction of signaling within platelets that causes intraplatelet Ca^2+^ release and activation of the platelet integrin, α_IIb_β_3_.(Mazzucato et al., 2002, Nesbitt et al., 2002, Kasirer-Friede et al., 2004, Constantinescu-Bercu et al., 2020) Physiologically, these signaling events are highly dependent on flow, as once platelets are bound to VWF, these shear forces induce unfolding of the mechanosensitive juxtamembrane region of GPIbα that translates the mechanical signal into intracellular biochemical events.(Zhang et al., 2015, Ju et al., 2016) Signaling events through GPIbα are dependent upon the binding of adaptor and signaling molecules such as Src kinases, Lyn and c-Src, 14-3-3 isoforms and phosphoinositide-3 kinase (PI3K) that can associate with the GPIbα intracellular tail.(Gu et al., 1999, Mangin et al., 2004, Mangin et al., 2009, Dai et al., 2005, Mu et al., 2008) Downstream activation of PLCγ2, PI3K-Akt, cGMP-PKG, mitogen activated kinase and LIM kinase 1 pathways have also been reported.(Wu et al., 2001, Li et al., 2006, Li et al., 2003, Yin et al., 2008, Garcia et al., 2005, Estevez et al., 2013, Mangin et al., 2003) By comparison to other platelet agonists (e.g. collagen, thrombin, ADP, thromboxane A2), the signaling effects of GPIbα are considered weak. In isolation, VWF-GPIbα signaling, which we refer to this as platelet ‘priming’ as opposed to activation, does not induce appreciable degranulation.(Constantinescu-Bercu et al., 2020) Therefore, the contribution of this ‘priming’ to normal hemostasis remains unclear as, in this setting, the effects of the other platelet agonists have the potential to mask those of GPIbα. However, in scenarios where other platelet agonists are either absent or in low abundance (e.g. platelet recruitment to the endothelial surface, or for some of the innate immune cell functions of platelets), the effects of GPIbα signaling may become more prominent.(Constantinescu-Bercu et al., 2020)

GPVI is a collagen/fibrin receptor on the platelet surface that belongs to the immunoglobulin superfamily.(Rayes et al., 2019, Alshehri et al., 2015) GPVI non-covalently associates with Fc receptor γ-chain (FcRγ) that signals via immunoreceptor tyrosine-based activation motifs (ITAM) following ligand binding.(Tsuji et al., 1997) Collagen binding to platelets induces clustering of GPVI, which results in the phosphorylation of FcRγ by Src family kinases, Lyn and Fyn, that associate with the intracellular domain of GPVI.(Schmaier et al., 2009, Ezumi et al., 1998) This causes the recruitment and phosphorylation of Syk tyrosine kinase, and formation of a LAT-based signaling complex that can activate phospholipase C (PLC) γ2 and lead to release of intraplatelet Ca^2+^ stores, activation of protein kinase (PK) C, and ultimately α_IIb_β_3_ activation and both α- and dense-granule release.(Rayes et al., 2019)

Previous studies have suggested functional associations between GPIbα and GPVI and/or its co-receptor FcRγ.(Falati et al., 1999, Wu et al., 2001, Arthur et al., 2005) For example, VWF-GPIbα-mediated platelet responses are reportedly impaired in GPVI/FcRγ deficiencies in both mice and humans.(Wu et al., 2001, Goto et al., 2002) There is also evidence that VWF can potentiate responses after collagen mediated responses in human platelets.(Baker et al., 2004) However, the molecular basis of GPIbα and GPVI receptor crosstalk has not been elucidated. Using a novel GPIbα transgenic mouse in which the last 24 amino acids (a.a.) of the GPIbα intracellular tail were deleted, we demonstrate the importance of this region not only to VWF-dependent signaling in platelets, but also reveal a major contribution in augmenting GPVI-mediated platelet signaling, further underscoring the importance of GPIbα in thrombus formation, beyond its well-described role in VWF-mediated platelet recruitment.

## Results

### Generation of GpIbα^Δsig/Δsig^ mice

There is a very high sequence identity between the intracellular tails (which harbors the filamin, 14-3-3 and PI3K binding regions) of human and murine GPIbα, supporting the contention that their functions are also well conserved (Figure 1A). To evaluate the influence of the GPIbα intracellular tail upon both VWF- and collagen/GPVI-mediated signaling, we generated a novel transgenic mouse (*GpIbα*^Δsig/Δsig^) using CRISPR-Cas9 technology. Using this approach, we introduced a point mutation (Ser695Stop) that resulted in a premature stop codon that deletes the last 24 amino acids of the GPIbα intracellular tail (a.a. 695-718) containing the entire 14-3-3 isoform and PI3K binding region, (Mangin et al., 2009, Mu et al., 2008) but maintains the upstream filamin binding site in GPIbα (residues 560-573 in human GPIbα; residues 668-681 in murine GPIbα)(Nakamura et al., 2006) (Figure 1A). Introduction of this premature stop codon in the GPIbα intracellular tail ensured normal transcriptional control of the *GpIbα* gene and maintained the extracellular and transmembrane domains (important for VWF binding function and association with GPIbβ), as well as the intracellular region that interacts with filamin. The introduction of the mutation was confirmed by genomic DNA sequencing (Figure 1B-C) and by Western blotting using an anti-GPIbα antibody that recognizes the terminal region of the intracellular tail (Figure 1D-E). Homozygous *GpIbα*^Δsig/Δsig^ mice were generated by mating mice heterozygous for the mutation, were born with the expected Mendelian frequencies and were both viable and fertile.

**Figure 1.**
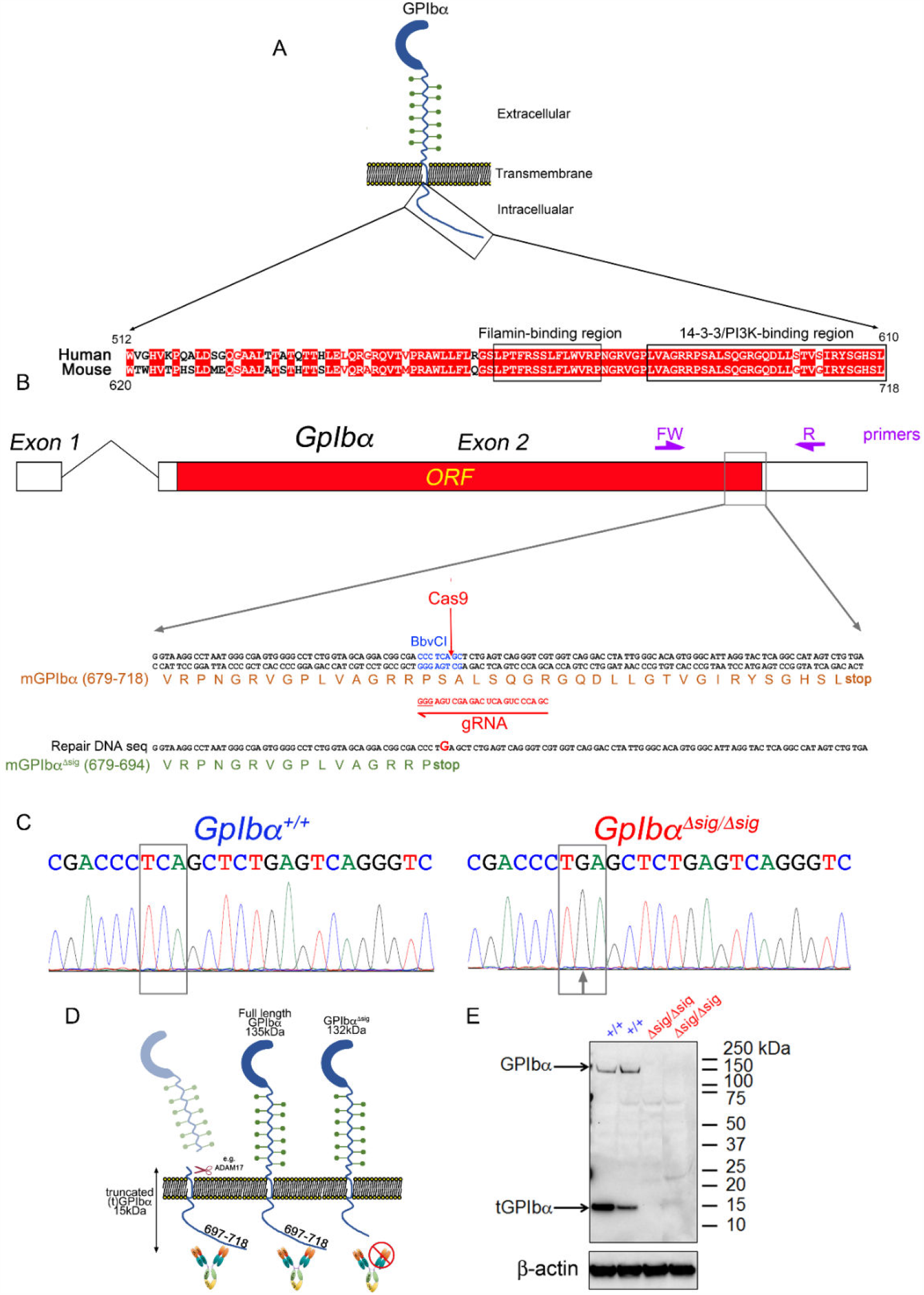
Generation and characterization of *GpIbα*^Δsig/Δsig^ mice. (A) Sequence alignment of the last 100 amino acids (a.a.) of human and mouse GPIbα. Sequence identities are highlighted in red. Filamin binding region: (a.a. 560-573) and (a.a. 668-681) for human and mouse GPIbα; PI3K/14-3-3 binding region: (a.a. 580-610) and (a.a. 688-718) for human and mouse GPIbα. (B) Schematic representation of the *GpIbα* gene with CRISPR guide target site, gRNA sequence, BbvCI restriction enzyme site and Cas9 predicted cut site. Primers used to amplify the *GpIbα* allele from genomic DNA are indicated in purple. Design of the 101 bp ssDNA repair template with the point mutation to introduce a codon stop eliminating the BbvCI restriction enzyme site and removing the last 24 a.a. of GPIbα is also shown. The resulting truncated amino acid sequence from *GpIbα*^Δsig/Δsig^ mice is indicated in green. (C) Genomic DNA sequences from *GpIbα*^+/+^ and *GpIbα*^Δsig/Δsig^ mice. Successful substitution is indicated with an arrow. (D) Diagram showing the binding of the anti-GPIbα tail Ab (Biorbyt; orb 215471). (E) Platelet lysates from *GpIbα*^+/+^ and *GpIbα*^Δsig/Δsig^ mice were probed with the anti-GPIbα tail and β-actin antibodies. Absence of band in the GPIbα western-blot confirms the successful truncation of the GPIbα intracellular tail in *GpIbα*^Δsig/Δsig^ mice.

### GpIbα^Δsig/Δsig^ mice platelet count, platelet size and hemostatic function

Hematological parameters in *GpIbα*^Δsig/Δsig^ mice were largely unaffected, with normal red blood cell and white blood cell counts, hemoglobin and hematocrit levels (Table S1). Complete *GpIbα* deficiency in mice causes severe thrombocytopenia and giant platelets, similar to the phenotype of Bernard-Soulier patients.(Ware et al., 2000, Lanza, 2006) However, *GpIbα*^*Δsig/Δsig*^ mice exhibited only a mild reduction (∼20%) in platelet count (Figure 2A). This reduction in platelet numbers in *GpIbα*^*Δsig/Δsig*^ mice was accompanied with slightly larger platelet size, as measured by flow cytometry (Figure 2B). Of note, the increase in platelet size was very modest and minimal when compared to the giant platelets associated with *GpIbα*^*-/-*^ mice.(Ware et al., 2000) Importantly, the expression of the major platelet receptors, GPVI, α_IIb_β_3_, GPIbβ, and the extracellular region of GPIbα was unaltered on *GpIbα*^*Δsig/Δsig*^ platelet surfaces (Figure 2C).

**Figure 2.**
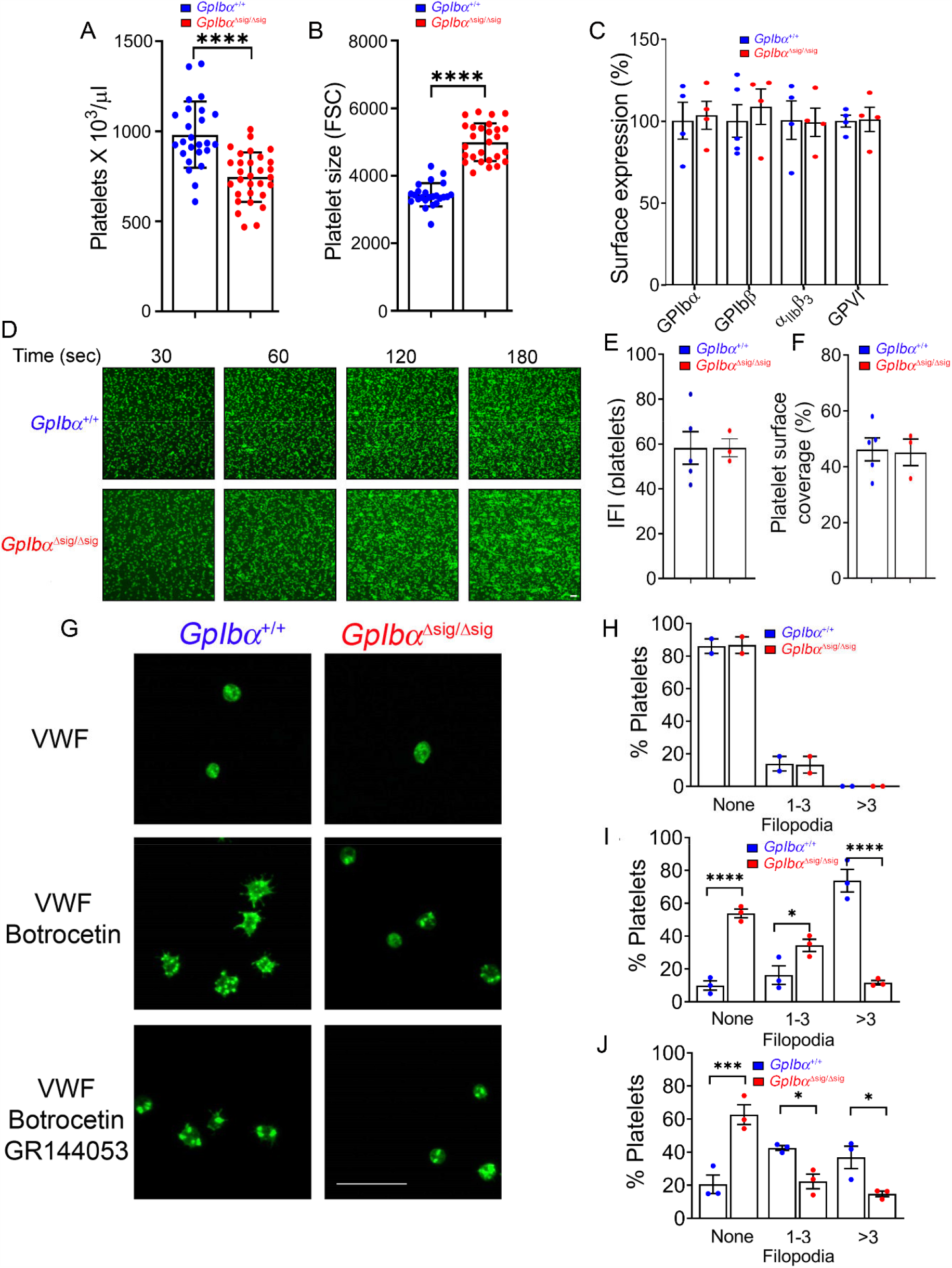
*GpIbα*^*Δsig/Δsig*^ platelets exhibit normal binding to VWF but disrupted GPIbα-mediated signaling. (A) Platelet counts and (B) platelet size in *GpIbα*^*+/+*^ (n=25) and *GpIbα*^*Δsig/Δsig*^ mice (n=30) as determined by flow cytometry. (C) Surface expression of platelet receptors GPIbα, GPIbβ, α_IIb_β_3_ and GPVI in *GpIbα*^*+/+*^ and *GpIbα*^*Δsig/Δsig*^ mice (n=4 for each genotype) determined by flow cytometry and expressed as % of control. (D-F) Plasma-free blood from *GpIbα*^*+/+*^ and *GpIbα*^*Δsig/Δsig*^ mice supplemented with anti-GPIbb-DyLight488 Ab was perfused over murine VWF at a shear rate of 1000s^-1^. (D) Representative fluorescence images (n≥3; scale bar 10 µm) and bar graphs analyzing the integrated fluorescence intensity (IFI) (E) and the surface coverage (F) of *GpIbα*^*+/+*^ and *GpIbα*^*Δsig/Δsig*^ platelets captured by murine VWF after 3.5 mins of flow. (G) Representative confocal images (n=3) of *GpIbα*^*+/+*^ and *GpIbα*^*Δsig/Δsig*^ platelets spread on mVWF and stained with Phalloidin-Alexa 488, in the absence or presence of Botrocetin or Botrocetin and GR144053 (scale bar 10 µm). (H-J) Percentage of platelets from *GpIbα*^*+/+*^ and *GpIbα*^*Δsig/Δsig*^ mice (n=2-3 for each genotype with individual data points representing the average of 3 fields of view) with no filopodia, 1-3 filopodia or >3 filopodia formed on murine VWF in the absence (H) or presence of Botrocetin (I), or Botrocetin and GR144053 (J). Data represents mean ± SEM was analyzed using unpaired two-tailed Student’s t-test (A-C;E-F) or using two-way ANOVA followed by Sidak’s multiple comparison test (H-J); *p<0.05, ***p<0.001, ****p<0.0001. Also see Figure S1 and Video 2.

To assess hemostatic function in *GpIbα*^*Δsig/Δsig*^ mice, we performed tail bleeding assays. Unlike *Vwf*^*-/-*^ mice or mice lacking the extracellular domains of GPIbα,(Denis et al., 1998, Ware et al., 2000, Kanaji et al., 2002) *GpIbα*^*Δsig/Δsig*^ mice displayed normal blood loss following tail transection (Figure 3A), suggesting that *GpIbα*^*Δsig/Δsig*^ platelets can be recruited to sites of vessel damage similar to wild-type mice.

**Figure 3.**
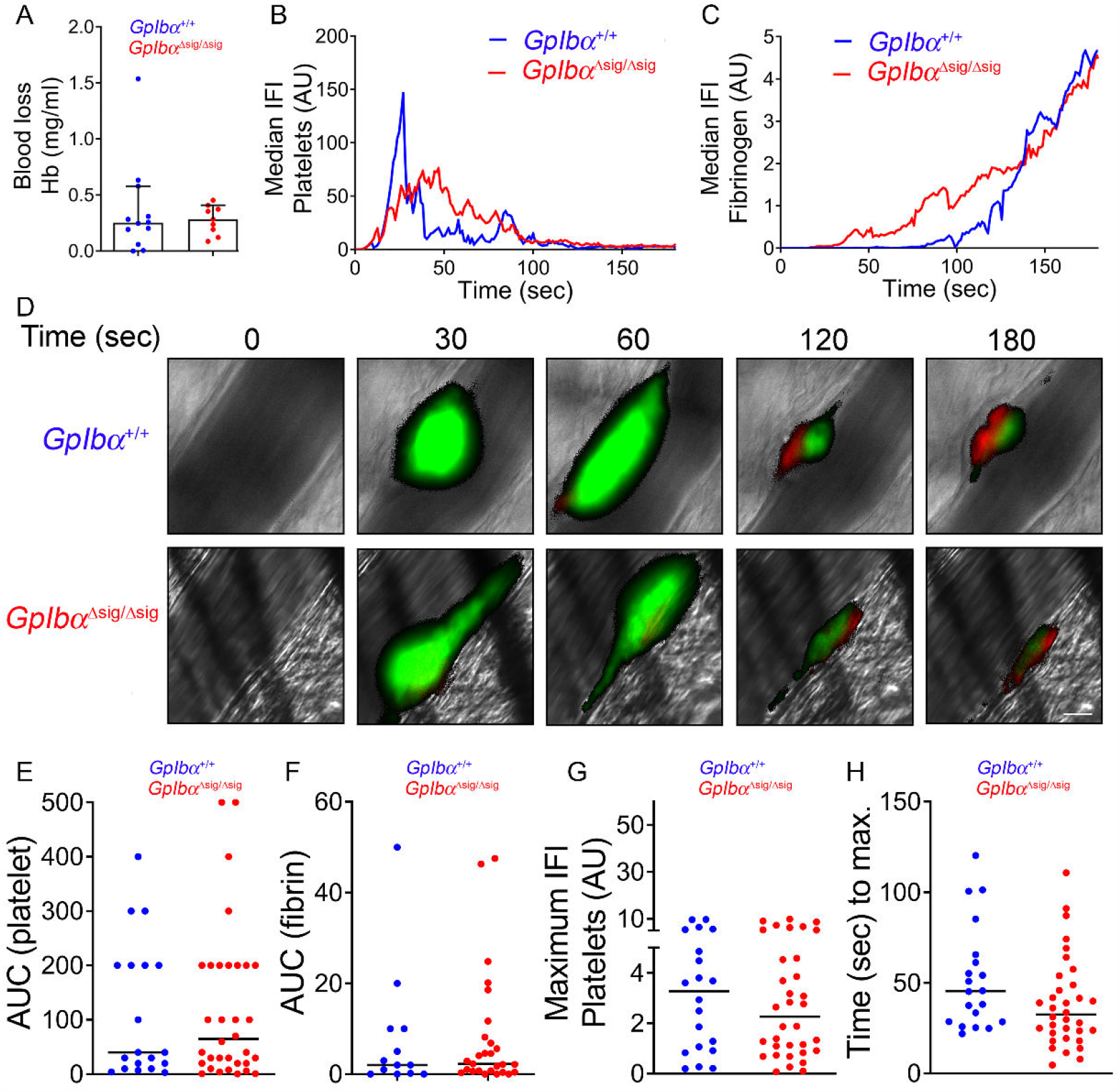
*GpIbα*^Δsig/Δsig^ mice display normal bleeding loss and platelet and fibrin accumulation in the laser-induced thrombosis model. (A) Bar graph analyzing blood loss after 10 min following tail transection in *GpIbα*^*+/+*^ and *GpIbα*^*Δsig/Δsig*^ mice (n=9 for each genotype). (B-H) Mice cremaster muscle arterioles were subjected to the laser-induced thrombosis model as described in SI Appendix, Methods. Curves represent median integrated fluorescence intensity (IFI) from platelets (arbitrary units: AU) (B) or fibrin(ogen) (C) as a function of time after the injury (20 thrombi in 3 *GpIbα*^*+/+*^ and 34 thrombi in 4 *GpIbα*^*Δsig/Δsig*^ mice). (D) Representative composite fluorescence images of platelets (green) and fibrin (red) with bright field images after laser-induced injury of the endothelium of *GpIbα*^*+/+*^ (top panels) versus *GpIbα*^*Δsig/Δsig*^ mice (bottom panels). Scale bar represents 10 µm. Graphs showing the area under curve values from the platelet IFI (E) or fibrin(ogen) IFI (F) vs time from individual thrombus. (G) Distribution of the maximal thrombus size expressed in IFI platelet arbitrary units (AU) and (H) the time to maximal thrombus size. Each symbol represents one thrombus. Horizontal lines intersecting the data set represent the median. Data was analyzed using Mann Whitney test; ns: p>0.05. Also see Video 1.

Next, we analyzed thrombus formation in *GpIbα*^*Δsig/Δsig*^ mice following mild laser injury to cremaster arteriole vascular wall.(Stalker, 2020, Salles-Crawley et al., 2014, Crawley et al., 2019) In this model, there was no difference between *GpIbα*^*Δsig/Δsig*^ mice and wild-type littermates in thrombus formation, as measured by the kinetics and extent, of both platelet accumulation and fibrin deposition (Figure 3B-H; Video 1). These results support the contention that deletion of the GPIbα does not appreciably influence either platelet recruitment or their ability to support thrombin generation. Importantly, platelet accumulation following mild laser injury requires both VWF and thrombin but has less dependency upon collagen exposure or GPVI signaling. (Dubois et al., 2007, Dubois et al., 2006) Collectively, our data indicate that although the major filamin binding site in GPIbα was preserved, deletion of the last 24 a.a. modestly influences platelet size and numbers. However, despite this, lack of the terminal portion of the intracellular tail in *GpIbα*^*Δsig/Δsig*^ mice does not alter the hemostatic response to tail-transection or in laser-induced thrombosis.

### GpIbα^Δsig/Δsig^ platelets bind VWF normally, but exhibit decreased VWF-mediated signaling

To more specifically examine the effect of the GPIbα intracellular tail truncation upon VWF-dependent platelet capture, we used microchannels coated with murine VWF over which we perfused plasma-free blood (to remove fibrinogen and outside-in activation α_IIb_β_3_) at 1000s^-1^. Under these conditions, *GpIbα*^*Δsig/Δsig*^ platelets were recruited normally to murine VWF-coated surfaces (Figure 2D; Video 2). This was quantified by measuring both surface coverage and platelet accumulation (measured by total platelet fluorescence), both of which were unaltered when compared to plasma-free blood *GpIbα*^*+/+*^ littermates (Figure 2E-F). In line with the normal expression of the extracellular domain of GPIbα, and also consistent with the *in vivo* data, these results more formally demonstrate that *GpIbα*^*Δsig/Δsig*^ platelets are recruited normally to VWF surfaces under flow.

To investigate the impact of the deletion of the last 24 a.a. GPIbα on downstream VWF signaling responses, we performed platelet spreading assays on murine VWF surfaces, which rely upon VWF-GPIbα signaling. Under basal static conditions, very few platelets (either *GpIbα*^*+/+*^ or *GpIbα*^*Δsig/Δsig*^) bound to VWF. However, of those that did, very few exhibited filopodia (13.9%±4.5 and 13.2%±5.2, respectively) (Figure 2G and 2H). When these experiments were repeated in the presence of botrocetin, a large proportion (90%±2.8) of *GpIbα*^*+/+*^ platelets underwent shape changes and developed filopodia, a well-described consequence of VWF-GPIbα signaling (Figure 2G and 2I; Figure S1A-B).(Mangin et al., 2004, Mangin et al., 2003) When similar experiments were performed in the presence of both botrocetin and GR144053, which competitively inhibits the interaction of α_IIb_β_3_ with VWF and/or fibrinogen (Figure 2G and 2J), the number of *GpIbα*^*+/+*^ platelets forming filopodia was not appreciably influenced (Figure S1B), but the proportion of that formed >3 filopodia was significantly reduced (37%±6.7 vs. 74%±6.9) (Figure S1A), revealing the contribution of outside-in signaling to filopodia formation.

Strikingly, although *GpIbα*^*Δsig/Δsig*^ platelets bound VWF surfaces in the presence of botrocetin, they had a significantly diminished ability to form filopodia (46%±2.6) when compared to *GpIbα*^*+/+*^ platelets (90%±2.8) (Figure 2G and 2I; Figure S1C-D). This difference in filopodia formation was also observed in the presence of GR144053 (Figure 2G and 2J). Moreover, GR144053 had no effect upon filopodia formation in *GpIbα*^*Δsig/Δsig*^ platelets (Figure S1C), suggesting that the reduced filopodia formation in these platelets was likely due to a defect in VWF-GPIbα signaling manifest by a lack of activation of α_IIb_β_3_ in response to VWF-GPIbα binding. Taken together, these results indicate that whereas deletion of the last 24 a.a. of the intracellular tail of GPIbα does not influence platelet binding to VWF, it significantly reduces VWF-GPIbα downstream signaling response including α_IIb_β_3_ activation.

### The intracellular tail of GPIbα is important for GPVI signaling

Platelet function in *GpIbα*^*Δsig/Δsig*^ mice was further evaluated by analysis of α_IIb_β_3_ integrin activation and P-selectin exposure in response to different agonists using flow cytometry, and also by aggregometry. In response to increasing concentrations of ADP, washed *GpIbα*^*Δsig/Δsig*^ platelets exhibited normal α_IIb_β_3_ activation and P-selectin exposure (Figure 4A-B). Platelet aggregation in response to ADP was also normal (Figure 4C-D). Responses to thrombin were also normal except for a slight, but significant, decrease in P-selectin exposure (and a non-significant decrease in activation of α_IIb_β_3_) following stimulation with the lowest thrombin concentration (Figure 4A-B). Despite this, platelet aggregation in response to thrombin was normal (Figure 4C-D). The cause of the reduced P-selectin exposure in response to low thrombin concentration is unclear, but this may reflect the findings of a previous study that suggested the importance of 14-3-3ζ binding to GPIbα specifically for low-dose thrombin responses.(Estevez et al., 2016)

**Figure 4:**
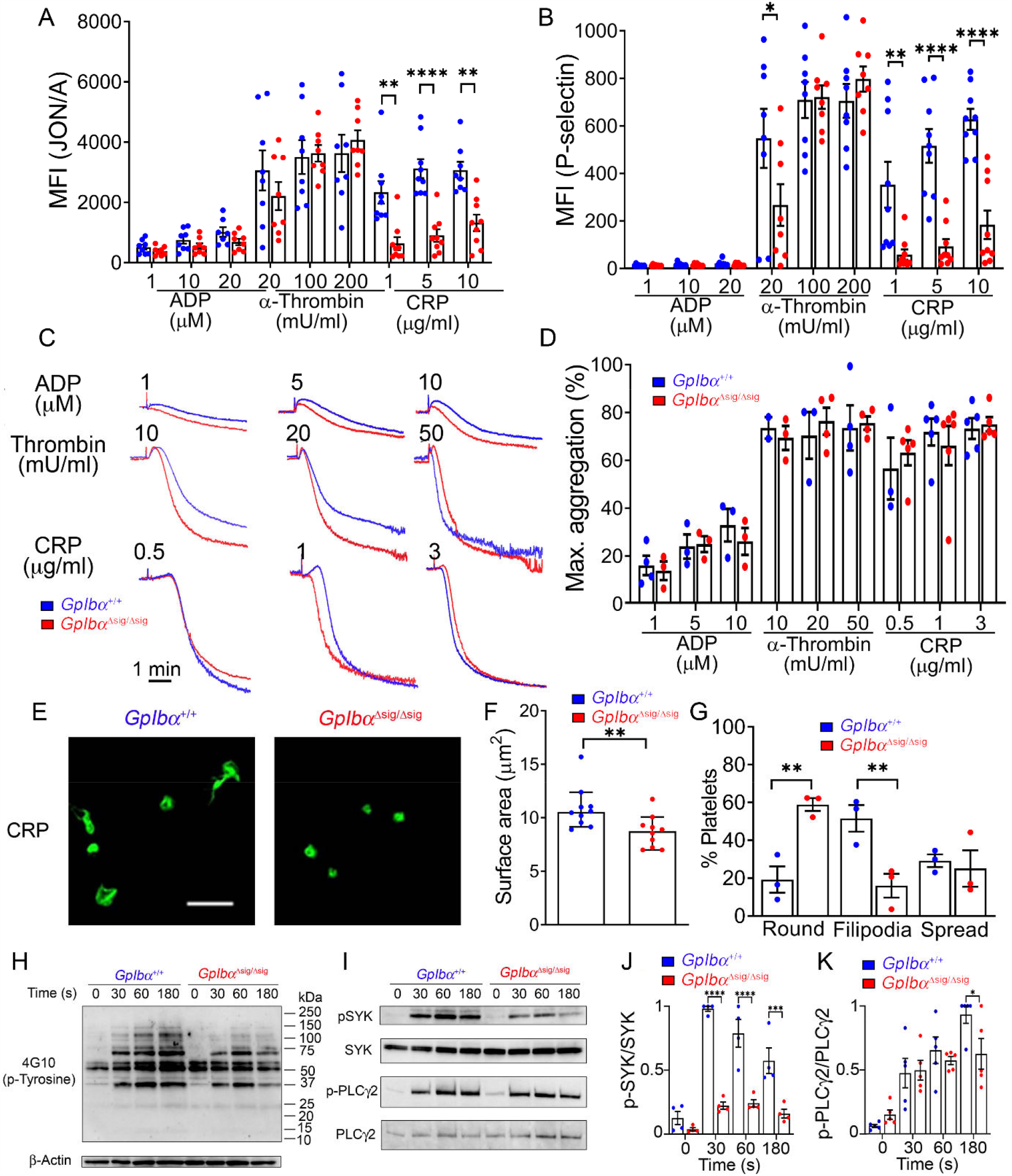
*GpIbα*^*Δsig/Δsig*^ platelets exhibit altered GPVI-mediated signaling. **(A-B)** Flow cytometric analysis of surface expression of activated α_IIb_β_3_ (**A**) and P-selectin (**B**) in *GpIbα*^*+/+*^ and *GpIbα*^*Δsig/Δsig*^ platelets (n=8) in response to ADP (1-20μM), α-thrombin (20-200mU/ml), or CRP (1-10μg/ml). MFI: geometric mean fluorescence intensity **(C)** Representative aggregation traces (n=3-6) of washed platelets isolated from *GpIbα*^*+/+*^ (blue) or *GpIbα*^*Δsig/Δsig*^ (red) mice and stimulated with ADP (1-10μM), α-thrombin (20-50mU/ml) or CRP (0.5-3μg/ml). Aggregation was monitored using a Chronolog aggregometer over 6 mins. **(D)** Bar graph analysing the maximum aggregation (%) obtained in the conditions presented in (**C**). **(E)** Representative micrographs (n=3 for each genotype; 3 fields of view analyzed per condition; scale bar 10 µm) of *GpIbα*^*+/+*^ and *GpIbα*^*Δsig/Δsig*^ platelets (>400) spread on CRP and stained with Phalloidin-Alexa 488. Bar graphs quantifying the surface area (**F)** and percentages (**G**) of platelets that remained round, formed filopodia or spread on CRP. **(H)** Western blot analyzing tyrosine kinase phosphorylation in platelets from *GpIbα*^*+/+*^ and *GpIbα*^*Δsig/Δsig*^ mice, following stimulation with 3 µg/ml CRP for 0-180s, using β-actin as a loading control (representative of n=3). **(I)** Western blots analyzing the levels of phosphorylated and non-phosphorylated SYK and PLCγ2 in platelets from GpIbα^+/+^ and GpIbα^Δsig/Δsig^ mice, after 0-180s stimulation with CRP (representative of n=3). **(J-K)** Bar graphs displaying the levels of phosphorylated SYK and PLCγ2 in platelets from GpIbα^+/+^ and GpIbα^Δsig/Δsig^ mice, after 0-180s stimulation with CRP and normalizing the intensity according to the non-phosphorylated levels of SYK and PLCγ2. For the surface area (F), the data represent the median±CI and was analyzed using the unpaired Mann Whitney test. All other data is displayed as mean±SEM and was analyzed using two-way ANOVA followed by Sidak’s multiple comparison test. *p<0.05, **p<0.01, ***p<0.001, ****p<0.0001. Also see Figure S2 and S3.

Despite largely unaffected responses to ADP and thrombin, in response to all tested concentrations (1-10μg/ml) of collagen-related peptide (CRP), *GpIbα*^*Δsig/Δsig*^ platelets exhibited markedly reduced α_IIb_β_3_ activation and P-selectin exposure (Figure 4A-B). Importantly, platelets still exhibited a concentration-dependent response to CRP, suggesting that signaling had been dampened, rather than ablated. Interestingly, *GpIbα*^*Δsig/Δsig*^ platelet aggregation following CRP stimulation appeared normal (Figure 4C-D).

Next, we evaluated the ability of *GpIbα*^*Δsig/Δsig*^ platelets to spread on fibrinogen surfaces with and without prior stimulation with thrombin. Without platelet stimulation, similar to wild-type platelets, most *GpIbα*^*Δsig/Δsig*^ platelets remained round, and very few exhibited filopodia or spread (FigureS2A and S2D). When wild-type or *GpIbα*^*Δsig/Δsig*^ platelets were activated with thrombin, ∼80% platelets of both genotypes spread fully (Figure S2B and S2E), with no difference observed in the spread platelet area (Figure S3C). As full spreading is highly dependent upon outside-in signaling through α_IIb_β_3_,(Durrant et al., 2017) this suggests that this signaling pathway is unaffected in *GpIbα*^*Δsig/Δsig*^ platelets. We then explored the ability of platelets to spread on CRP-coated surfaces. Consistent with diminished platelet activation in response to CRP, 59%±3.4 of *GpIbα*^*Δsig/Δsig*^ platelets remained round in contrast to only 19%±7 of wild-type platelets (Figure 4E and 4G). This effect was also quantified by a 20% reduction in bound platelet area (Figure 4F) and in the reduced incidence of filopodia formation - 16%±6.3 for *GpIbα*^*Δsig/Δsig*^ vs 52%±7.1 for *GpIbα*^*+/+*^ (Figure 4E and 4G). Collectively, these results reveal an appreciable defect in GPVI-mediated signaling in *GpIbα*^*Δsig/Δsig*^ platelets.

To explore the mechanism underlying the defect in GPVI-mediated signaling, we assessed whole-cell tyrosine phosphorylation (p-Tyr) over time in *GpIbα*^*+/+*^ and *GpIbα*^*Δsig/Δsig*^ platelets after stimulation with 3µg/ml CRP. There was an overall reduction in tyrosine phosphorylation in *GpIbα*^*Δsig/Δsig*^ platelets compared to wild-type platelets (Figure 4H). Further analysis revealed appreciably reduced Syk kinase activation in *GpIbα*^*Δsig/Δsig*^ platelets, as measured by phosphorylation of Syk on Tyr525 and Tyr526, in response to CRP (Figure 4I-J). There was also some evidence of lower phosphorylation levels of one of its downstream targets PLCγ2 (at Tyr1217), although this was less marked than for those observed with pSyk (Figure 4I and 4K).

To assess whether the effect of truncation of GPIbα was specific for GPVI-mediated platelet responses, or whether other tyrosine-mediated signaling pathways might also be affected, we stimulated *GpIbα*^*Δsig/Δsig*^ and wild-type platelets with rhodocytin (C-type lectin receptor 2 (CLEC-2) agonist). In these experiments, the tyrosine-phosphorylation profile of *GpIbα*^*Δsig/Δsig*^ platelets in response to rhodocytin was very similar to that of *GpIbα*^*+/+*^ platelets (Figure S3A) with only slightly reduced phosphorylation of Syk (Figure S3B-C). Overall, α_IIb_β_3_ activation and P-selectin exposure in response to rhodocytin was unaffected (Figure S3D-E), suggesting that the GPIbα tail is not of major importance for CLEC-2 ITAM-mediated signaling.

### The role of the GPIbα intracellular tail in platelet recruitment and aggregation under flow

To examine the consequences of the combined effects of disrupted VWF-GPIbα signaling and diminished GPVI-signaling in platelets in a more physiological setting, we studied platelet recruitment and aggregate formation on collagen-coated microchannels using an *ex vivo* microfluidic assay. Experiments were performed at high (3000s^-1^), medium (1000s^-1^) and low (200s^-1^) shear, as platelet <u>recruitment</u> is increasingly dependent on VWF-GPIbα as shear increases. Once recruited, subsequent platelet aggregation and accumulation on collagen surfaces becomes more dependent upon GPVI signaling (Ruggeri, 2007, Kuijpers et al., 2004, Nieswandt et al., 2001).

Perfusing whole blood at high shear (3000s^-1^) over collagen, we observed a marked reduction in surface coverage (3.6±1.2%) of *GpIbα*^*Δsig/Δsig*^ platelets after 3 mins when compared to *GpIbα*^*+/+*^ platelets (27.4±4.7%) (Figure 5A-B; Video 3). At high shear rates, initial surface coverage is highly dependent upon VWF-mediated platelet recruitment. The deficit in surface coverage of *GpIbα*^*Δsig/Δsig*^ platelets on collagen suggests that VWF-GPIbα signaling and therefore early/rapid activation of cell surface α_IIb_β_3_ may be important for stabilizing platelet recruitment at higher shear rates. *GpIbα*^*Δsig/Δsig*^ platelets that bound to collagen also formed smaller aggregates (measured by total platelet fluorescence) than those from *GpIbα*^*+/+*^mice (Figure 5C), likely reflecting the subsequent effect of diminished collagen-GPVI signaling.

**Figure 5.**
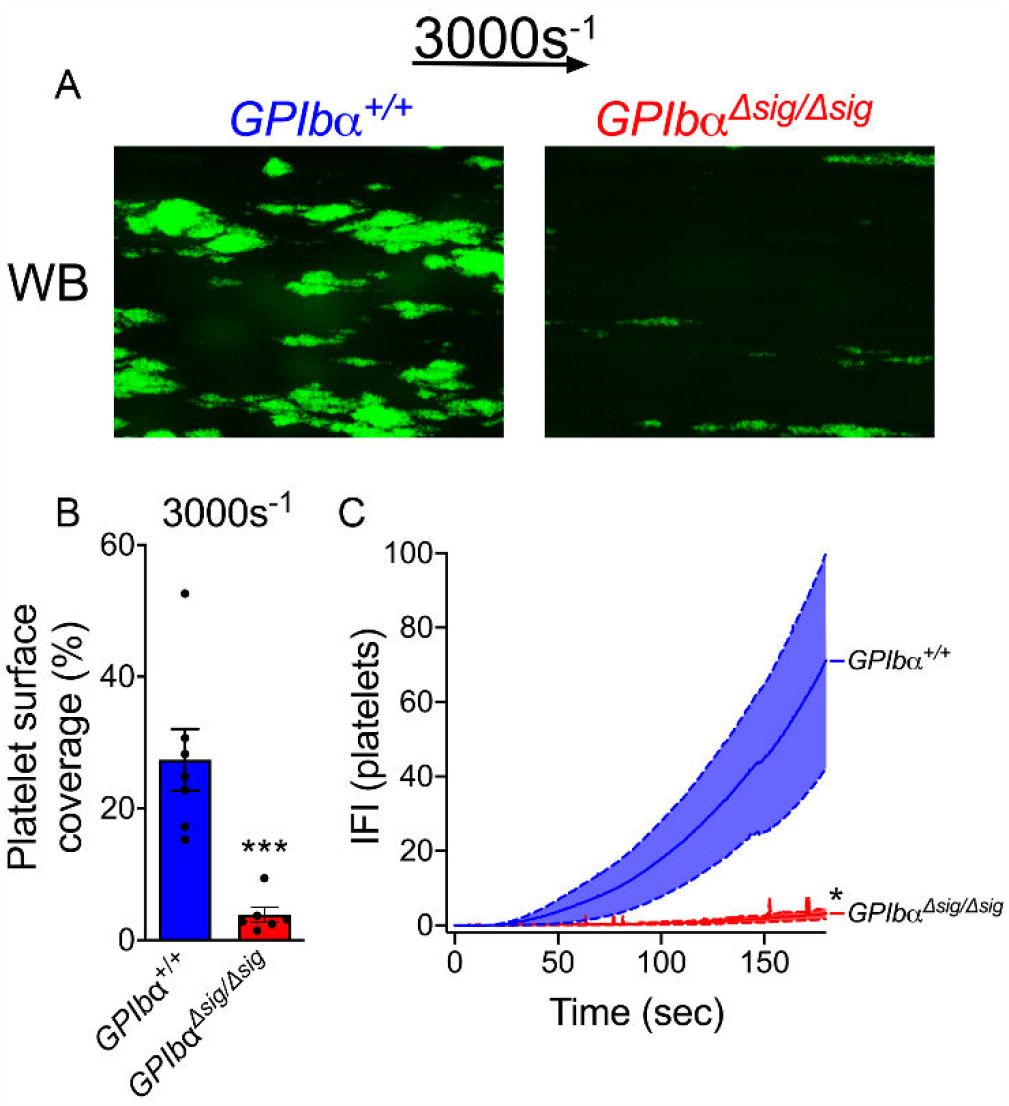
*GpIbα*^*Δsig/Δsig*^ platelets have a reduced ability to bind to collagen and form microthrombi at 3000s^-1^. Hirudin anticoagulated whole blood from *GpIbα*^+/+^ and *GpIbα*^Δsig/Δsig^ mice was labelled with anti-GPIbβ-DyLight488 Ab and perfused over fibrillar collagen type I (0.2 mg/ml) at a shear rate of 3000s^-1^ for 3 mins. **(A)** Representative fluorescence images (n=6) after 3 minutes of perfusion in whole blood (WB) from *GpIbα*^+/+^ and *GpIbα*^Δsig/Δsig^ mice at 3000s^-1^. Platelet deposition **(B)** and thrombus build-up measured as integrated fluorescence intensity (IFI) (**C**). All data is shown as mean ± SEM and analyzed using unpaired two-tailed student’s t-test The maximal platelet IFI was used to compare the thrombus build up data. *p<0.05, ***p<0.001. Scale bar 100 µm. Also see Video 3.

Experiments at high shear are limited by the quantity of mouse blood required for each experiment, which restricted further analyses at 3000s^-1^. However, similar results were also observed on collagen-coated microchannels at 1000s^-1^ shear rates. Both platelet surface coverage (Figure 6A-B) and platelet accumulation (Figure 6A and 6C) were markedly reduced using blood from *GpIbα*^*Δsig/Δsig*^ mice when compared to *GpIbα*^*+/+*^ mice (Video 4).

**Figure 6:**
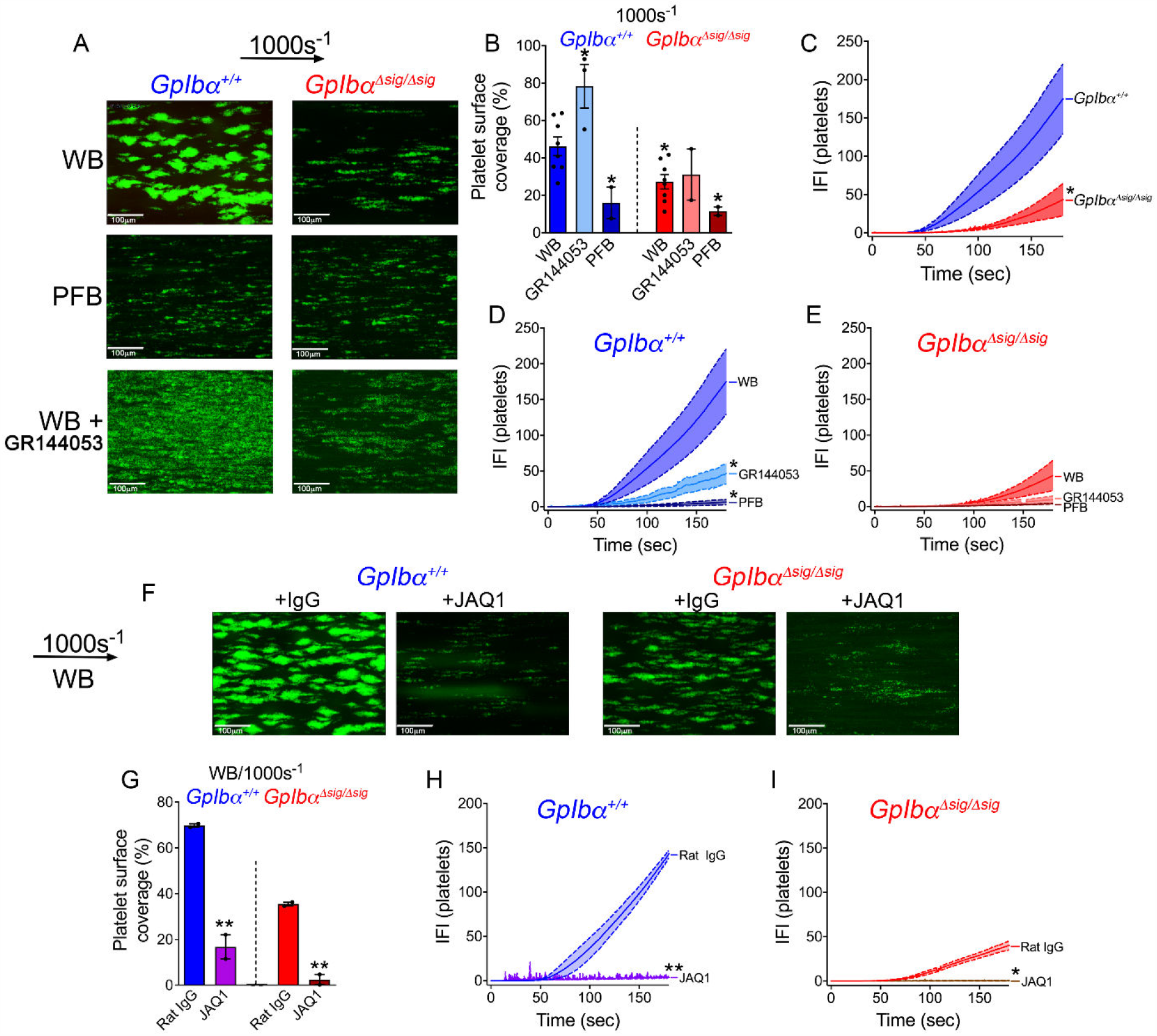
*GpIbα*^*Δsig/Δsig*^ platelets have a reduced ability to bind to collagen and form microthrombi at 1000s^-1^. **(A-E)** Hirudin anticoagulated whole blood supplemented or not with GR144053 or plasma-free blood from *GpIbα*^+/+^ and *GpIbα*^Δsig/Δsig^ mice was labelled with anti-GPIbβ-DyLight488 Ab and perfused over fibrillar collagen type I (0.2 mg/ml) at a shear rate of 1000s^-1^ for 3 mins. (**A**) Representative fluorescence images (n=2-4) after 3 minutes of perfusion in whole blood (WB), plasma-free blood (PFB) or WB + GR144053 from *GpIbα*^+/+^ and *GpIbα*^Δsig/Δsig^ mice. Platelet deposition **(B)** and thrombus build-up measured as IFI (**C-E**). All data is shown as mean ± SEM and analyzed using unpaired two-tailed student’s t-test (**C**) or one-way ANOVA followed by Dunnett’s multiple comparison test **(B, D-E)**. Data is compared to means from *GpIbα*^+/+^ WB (**B**,**D**) or *GpIbα*^Δsig/Δsig^ WB (**E**). The maximal platelet IFI was used to compare the thrombus build up data. *p<0.05. Scale bar 100 µm. Also see Video 4. **(F-I)** Hirudin anticoagulated whole blood from *GpIbα*^+/+^ and *GpIbα*^Δsig/Δsig^ mice supplemented with JAQ1 or Rat-IgG control Abs (20µg/ml) was labelled with anti-GPIbβ-DyLight488 Ab and perfused over fibrillar collagen type I (0.2 mg/ml) at a shear rate of 1000s^-1^ for 3 mins. **(F)** Representative fluorescence images (n=2-3) after 3 minutes of perfusion. Platelet deposition **(G)** and thrombus build-up measured as IFI (**H**,**I**). All data is shown as mean ± SEM and analyzed using unpaired two-tailed student’s t-test. The maximal platelet IFI was used to compare the thrombus build up data. *p<0.05, **p<0.01. Scale bar 100 µm.

When experiments were performed in plasma-free blood (to remove soluble VWF and fibrinogen), the surface coverage of *GpIbα*^*Δsig/Δsig*^ and *GpIbα*^*+/+*^ platelets on collagen (mediated through direct interactions of platelets with collagen) was reduced to similar levels (Figure 6A-B) showing that a small amount of VWF-independent binding to collagen occurs at 1000s^-1^. Both *GpIbα*^*Δsig/Δsig*^ and *GpIbα*^*+/+*^ platelet accumulation on collagen (measured by total platelet fluorescence) was limited in plasma-free conditions due to lack of VWF-mediated recruitment, but also due to the absence of fibrinogen necessary to enable platelet aggregation, which prevented platelet accumulation beyond the first layer of platelets (Figure 6A, 6D and 6E).

When whole blood experiments were performed in the presence of GR144053, to block α_IIb_β_3_, we observed an increase in surface coverage with wild-type platelets accompanied with an overall decrease in thrombus formation due to the formation of a platelet monolayer (Figure 6A-B).(Pugh et al., 2010) This effect is due to the inhibition of platelet aggregation that promotes the formation of focal microthrombi that reduces apparent surface coverage (Figure 6A-B). Interestingly, this effect of GR144053 on surface coverage was not observed with *GpIbα*^*Δsig/Δsig*^ platelets. Surface coverage as well as platelet accumulation of *GpIbα*^*Δsig/Δsig*^ platelets was similar in both the absence and presence of GR144053 (Figure 6A-B). This may reflect the lack of VWF-GPIbα signaling/VWF-dependent α_IIb_β_3_ activation combined with decreased collagen/GPVI signaling-mediated α_IIb_β_3_ activation in whole blood, which is why blocking α_IIb_β_3_ does not influence these parameters in *GpIbα*^*Δsig/Δsig*^ blood (Figure 6A-B and 6E).

To more specifically examine the role of GPVI in this system, we performed experiments in the presence of JAQ1, an anti-murine GPVI blocking antibody. Blocking GPVI had a significant effect upon both the surface coverage and platelet accumulation of both *GpIbα*^*Δsig/Δsig*^ and *GpIbα*^*+/+*^ platelets, revealing the important contribution of GPVI signaling at 1000s^-1^ (Figure 6F-I), both in stabilizing platelet recruitment and their subsequent aggregation.

We then examined these same processes at low, venous shear rates (200s^-1^) (Figure 7). Under these conditions, the dependencies on VWF and collagen were slightly different to those at 1000s^-1^. Surface coverage observed for *GpIbα*^*Δsig/Δsig*^ platelets was slightly reduced compared to *GpIbα*^*+/+*^ platelets although not reaching significance (Figure 7A-B), but thrombus growth was significantly diminished (Figure 6C; Video 5). Of note, overall platelet accumulation (Figure 7C) was appreciably lower when compared to higher shear conditions (Figure 6C), in part, due to the smaller volume of blood perfused through the channels at lower shear over the 3 min time frame of the experiment.

**Figure 7:**
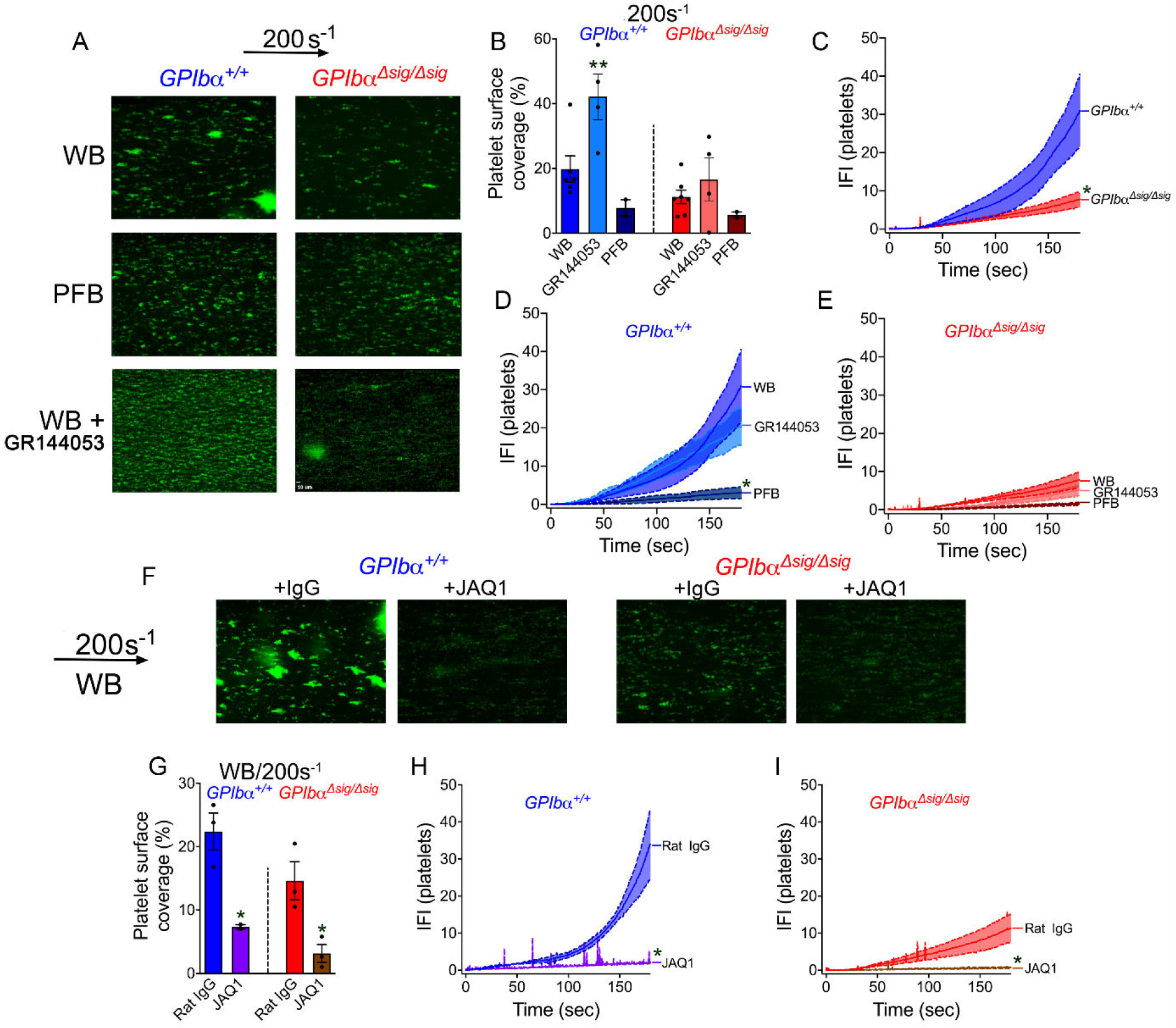
*GpIbα*^*Δsig/Δsig*^ platelets have a reduced ability to bind to collagen and form microthrombi at 200s^-1^. **(A-E)** Hirudin anticoagulated whole blood supplemented or not with GR144053 or plasma-free blood from *GpIbα*^+/+^ and *GpIbα*^Δsig/Δsig^ mice was labelled with anti-GPIbβ-DyLight488 Ab and perfused over fibrillar collagen type I (0.2 mg/ml) at a shear rate of 200s^-1^ for 3 mins. (**A**) Representative fluorescence images (n=2-4) after 3 minutes of perfusion in whole blood (WB), plasma-free blood (PFB) or WB + GR144053 from *GpIbα*^+/+^ and *GpIbα*^Δsig/Δsig^ mice. Platelet deposition **(B)** and thrombus build-up measured as IFI (**C-E**). All data is shown as mean ± SEM and analyzed using unpaired two-tailed student’s t-test (**C**) or one-way ANOVA followed by Dunnett’s multiple comparison test (**B**,**D-E**). Data is compared to means from *GpIbα*^+/+^ WB (**B**,**D**) or *GpIbα*^Δsig/Δsig^ WB (**E**). The maximal platelet IFI was used to compare the thrombus build up data. *p<0.05, **p<0.01. Scale bar 100 µm. Also see Video 5. **(F-I)** Hirudin anticoagulated whole blood from *GpIbα*^+/+^ and *GpIbα*^Δsig/Δsig^ mice supplemented with JAQ1 or Rat-IgG control Abs (20µg/ml) was labelled with anti-GPIbβ-DyLight488 Ab and perfused over fibrillar collagen type I (0.2 mg/ml) at a shear rate of 1000s^-1^ for 3 mins. **(F)** Representative fluorescence images (n=2-3) after 3 minutes of perfusion. Platelet deposition **(G)** and thrombus build-up measured as IFI (**H**,**I**). All data is shown as mean ± SEM and analyzed using unpaired two-tailed student’s t-test. The maximal platelet IFI was used to compare the thrombus build up data. *p<0.05. Scale bar 100 µm.

Using plasma-free blood, the surface coverage was similar for *GpIbα*^*Δsig/Δsig*^ and *GpIbα*^*+/+*^ platelets, mediated by direct (VWF-independent) interaction with collagen (Figure 7A-B). Similar to high-shear conditions, platelet accumulation under plasma-free conditions of *GpIbα*^*+/+*^ platelets was significantly reduced compared to whole blood (Figure 7D) similar to those observed with *GpIbα*^*Δsig/Δsig*^ blood (Figure 7E).

In the presence of GR144053, we saw the same increase in surface coverage of *GpIbα*^*+/+*^ platelets (Figure 6A-B) with reduced localized platelet build-up (Figure 7A) although the total platelet fluorescence/accumulation was not significantly different than in *GpIbα*^*+/+*^ whole blood (Figure 7D) likely due to the increased platelet coverage. Consistent with the results obtained under high-shear conditions, the effect of increased surface coverage in the presence of GR144053 was not observed with *GpIbα*^*Δsig/Δsig*^ platelets, nor was platelet accumulation appreciably further diminished (Figure 7A-B and 7E).

We next examined the role of GPVI in platelet aggregate formation at low shear using the GPVI-blocking JAQ1 antibody. With both *GpIbα*^*Δsig/Δsig*^ and *GpIbα*^*+/+*^ platelets, blocking GPVI significantly reduced surface coverage and platelet accumulation (Figure 7F-I). As removal of either VWF or blocking of GPVI had very similar effects, this suggests that VWF-GPIbα and GPVI-collagen binding may act synergistically to recruit platelets at low shear.

## Discussion

The ability of platelet GPIbα binding to VWF to transduce intraplatelet signaling events has been known for decades. However, the hemostatic role of this platelet ‘priming’ that follows has frequently been perceived as redundant due to the comparatively mild phenotypic changes in platelets that arise when compared to other platelet agonists (e.g. thrombin, collagen, ADP and thromboxane A2). Using a novel *GpIbα*^*Δsig/Δsig*^ mouse, we now demonstrate that the intracellular tail of GPIbα is important for transduction of both VWF-GPIbα signaling and collagen-GPVI-mediated responses in platelets (Figure 8).

**Figure 8:**
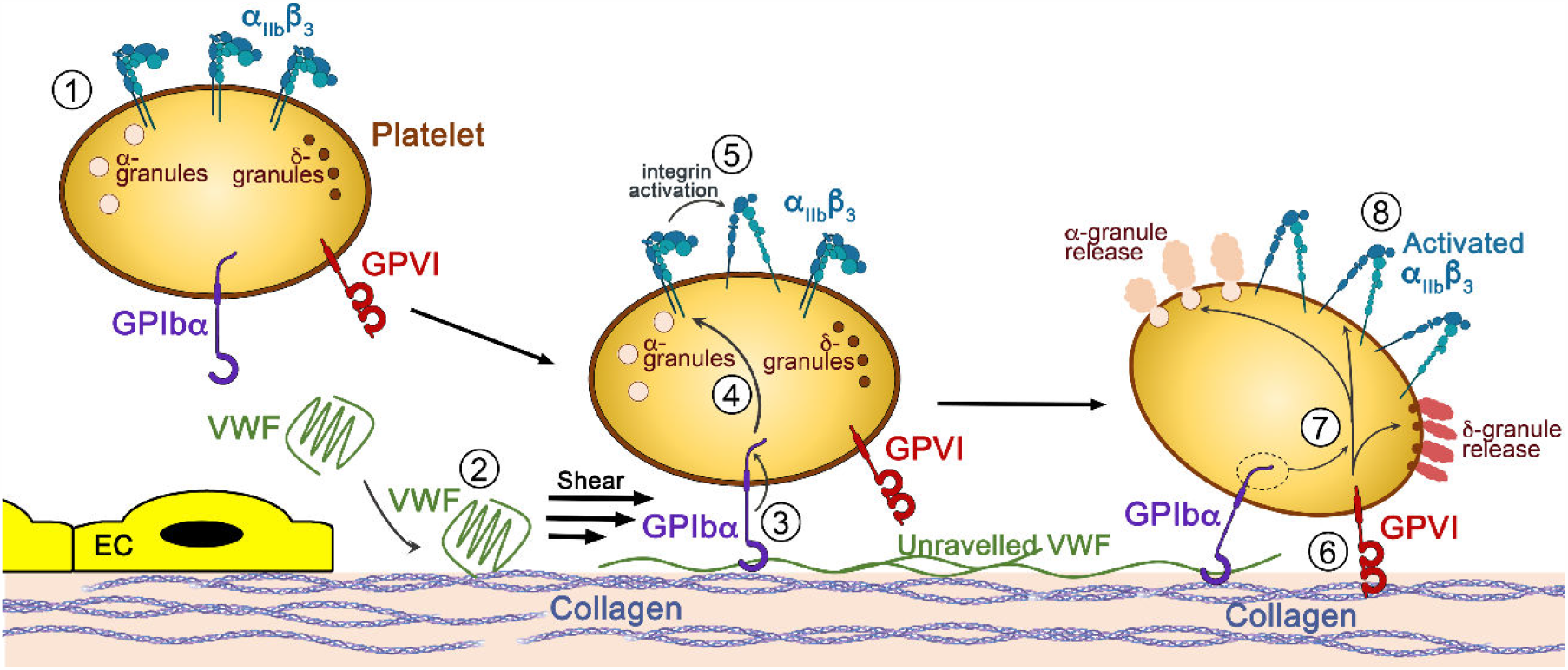
Proposed model for GPIbα-GPVI cross talk. Under normal conditions, resting/circulating platelets (1) present α_IIb_β_3_ on their surface in its closed conformation. Plasma VWF (2) circulates in its globular conformation with its A1 domain hidden, preventing interaction with platelet GPIbα. Upon vascular injury, the subendothelial extracellular matrix containing collagen becomes exposed to the blood. VWF, via its A3 domain, binds to collagen and, due to shear forces, unravels to expose its A1 domain to which platelet GPIbα binds (3). Next, mechanosensitive signaling events downstream of VWF A1-GPIbα that require the intracellular tail of GPIbα take place leading to some activation of surface α_IIb_β_3_ (4) while the deceleration of platelets allows for the subsequent binding of platelets to collagen via several collagen receptors including GPVI (6). The intracellular tail of GPIbα is also crucial for optimal collagen/GPVI signaling that lead to platelet activation, shape change and granule release (7). Ultimately, additional circulating platelets will be recruited at the site of injury to form the hemostatic plug (8).

The binding of GPIbα to VWF, and GPVI to collagen, are critical events for the formation of stable platelet aggregates under flow.(Ruggeri, 2007, Bergmeier et al., 2006, Nieswandt et al., 2011) Previous studies have suggested some interplay between these two signaling pathways. For example, physical associations between GPIbα and GPVI, or its co-receptor FcRγ, have been described.(Falati et al., 1999, Arthur et al., 2005) Moreover, VWF signaling has also been reported to potentiate platelet secretion in response to CRP.(Baker et al., 2004) Further crosstalk between these signaling pathways is also suggested by the altered VWF-GPIbα-dependent responses in human and mouse platelets deficient in GPVI signaling.(Wu et al., 2001, Goto et al., 2002)

To explore the signaling function of GPIbα and its influence upon GPVI signaling, we generated *GpIbα*^*Δsig/Δsig*^ mice by introducing an early stop codon by CRISPR/Cas9 technology to remove the last 24 amino acids GPIbα intracellular tail. The stop codon was introduced just downstream of the main filamin binding site (a.a. 668-681) (to maintain the role of filamin binding to normal platelet formation), but upstream of the 14-3-3 isoforms and PI3K binding regions that are important for VWF-GPIbα signaling.(Gu et al., 1999, Mangin et al., 2004, Mangin et al., 2009, Dai et al., 2005, Mu et al., 2008, Feng et al., 2003)(Figure 1). This approach resulted in uniform production of platelets that express GPIbα with truncated intracellular tail. This therefore circumvented many of the limitations associated with studying/expressing platelet receptor complexes in heterologous cellular systems to assay their function. Other mouse models of GPIbα have been generated including the full knockout (*GpIb*α^-/-^) and also *GpIb*α*/IL4R*α-tg mice that lack the extracellular region of GPIbα.(Ware et al., 2000, Kanaji et al., 2002) These two models do not enable analysis of VWF signaling, per se, as they lack the ability to bind VWF, meaning that one cannot dissociate the effects of loss of VWF binding and/or VWF signaling upon functional effects upon the platelets. The third mouse model is the hTg^Y605X^ transgenic mice that express human GPIbα that lacks the terminal 6 amino acids of the intracellular tail.(Kanaji et al., 2004) Although these mice have a phenotype associated with reduced megakaryocyte recovery following induced thrombocytopenia, more recent *in vitro* studies have revealed that these mice do not lack the entire 14-3-3/PI3K binding region,(Mu et al., 2008, Mangin et al., 2004, Mangin et al., 2009) suggesting that their VWF signaling function may not be fully disrupted making interpretation of the mouse phenotype difficult.

*GpIbα*^*Δsig/Δsig*^ mice had a modest reduction in platelet counts compared to *GpIbα*^*+/+*^ littermates (Figure 2A). The 20% decrease in *GpIbα*^*Δsig/Δsig*^ mouse platelet counts may likely be attributable to the small increase in platelet size (Figure 2B). Consistent with the flow cytometry results, we measured ∼16% increase in *GpIbα*^*Δsig/Δsig*^ platelet area compared to wild-type platelets when visualized by microscopy. Although the major filamin binding site remains intact, these findings may be consistent with CHO cell studies that suggested the presence of additional or extended filamin binding regions within intracellular tail of GPIbα.(Feng et al., 2003) Despite this, the effect upon platelet size is comparatively minor. Interestingly, platelet size is also moderately increased in the *GpIb*α*/IL4R*α-tg mice,(Kanaji et al., 2002) but, again, this is very modest compared to the size observed in *GpIbα*^*-/-*^ mice or in Bernard-Soulier platelets.(Ware et al., 2000, Lanza, 2006) It is worth mentioning that the 20% reduction in platelet count and slight increase in platelet size are unlikely to impart any discernible hemostatic defect.(Morowski et al., 2013)

The *GpIbα*^*Δsig/Δsig*^ mice exhibited normal hemostatic responses to tail transection, and normal thrombus formation following mild laser-induced thrombosis (Figure 3). As both of these models are sensitive to VWF function, these results are consistent with the normal VWF binding function of *GpIbα*^*Δsig/Δsig*^ platelets.(Denis et al., 1998, Dubois et al., 2007) Normal bleeding times were also reported in *hTg*^Y605X^ transgenic mice with no overt effect on platelet or coagulation functions.(Kanaji et al., 2004) Truncation of the intracellular tail of GPIbα did not alter expression of its extracellular domain (nor influence surface expression of GPIbβ, GPVI or α_IIb_β_3_) (Figure 2C). Consequently, *GpIbα*^*Δsig/Δsig*^ platelet capture to mouse VWF-coated surfaces was perhaps predictably unaffected (Figure 2D-F). Despite normal VWF binding though, deletion of the PI3K and 14-3-3 binding region in GPIbα (Mu et al., 2008, Mangin et al., 2004, Mangin et al., 2009) significantly diminished VWF-mediated signaling. *GpIbα*^*Δsig/Δsig*^ platelets exhibited markedly decreased filopodia extension upon stimulation of VWF binding with botrocetin (Figure 2G and 2I). Importantly, the same reduction in filopodia numbers in *GpIbα*^*Δsig/Δsig*^ platelets was observed in the presence of GR144053, an α_IIb_β_3_ antagonist, demonstrating that this effect is attributable to a defect in VWF-GPIbα signaling, as opposed to altered outside-in signaling induced by the VWF C4 domain binding to activated α_IIb_β_3_.(Figure 2G and 2J) This normal VWF-platelet capture observed in *GpIbα*^*Δsig/Δsig*^ mice is in line with previous studies showing that deletion of the 14-3-3ζ binding site in human GPIbα in GPIb-IX CHO cells does not affect VWF binding, but does reduce their ability to spread.(Mangin et al., 2004, David et al., 2010)

Demonstration of reduced platelet filopodia formation on VWF does not reveal the true physiological role VWF-mediated signaling. Other studies have endeavored to explore this using different approaches. For example, a membrane-permeable inhibitor of the 14-3-3ζ-GPIbα interaction (MP-αC) was shown to markedly inhibit GPIbα-dependent platelet agglutination (Dai et al., 2005) and was also protective in murine thrombosis models.(Yin et al., 2013) However, although this peptide disrupts the interaction between 14-3-3ζ and GPIbα, it may also influence 14-3-3ζ function independent of GPIbα binding. This contention is perhaps supported by a recent study revealing that whereas 14-3-3ζ deficient mice are protected against arterial thrombosis, these mice exhibit normal VWF-GPIbα-mediated platelet function.(Schoenwaelder et al., 2016) 14-3-3ζ must therefore be important for other GPIbα-independent platelet functions. Moreover, the binding and function of other 14-3-3 isoforms, which can also interact with the GPIbα intracellular tail, suggest that there may be some 14-3-3 redundancy, as far as GPIbα is concerned.(Schoenwaelder et al., 2016) The *GpIbα*^*Δsig/Δsig*^ mice now provide a more definitive means to examine the role of VWF-GPIbα signaling *in vivo*, particularly for some of the non-hemostatic functions of platelets that may be reliant upon this ‘priming’.

In addition to the defect in VWF-mediated signaling, *GpIbα*^*Δsig/Δsig*^ platelets exhibited markedly diminished collagen-mediated signaling through GPVI, supported by several lines of evidence. First, stimulation of *GpIbα*^*Δsig/Δsig*^ platelets with CRP resulted in significantly reduced surface expression of P-selectin and activation of α_IIb_β_3_ compared to *GpIbα*^*+/+*^ platelets (Figure 4A-B). Second, *GpIbα*^*Δsig/Δsig*^ platelets formed significantly fewer filopodia when spreading on CRP (Figure 4E-G). This effect was not due to a defect in outside-in signaling as *GpIbα*^*Δsig/Δsig*^ platelets adhered and spread normally on fibrinogen-coated surfaces both with and without thrombin stimulation (Figure S2). Third, the growth of *GpIbα*^*Δsig/Δsig*^ platelet thrombi on collagen under flow was severely reduced compared to wild-type platelets (Figs 5-7).

Bernard-Soulier patient platelets have historically been reported to respond normally to collagen (Andrews and Berndt, 2013). However, the thrombocytopenia and giant platelets associated with full GPIbα deficiency impair full analysis of other platelet signaling pathways under physiological flow conditions, which is also complicated by the loss of VWF-dependent platelet recruitment on collagen. It is worth noting though that the responsiveness of Bernard-Soulier platelets to collagen has been primarily characterized using platelet aggregation assays. Interestingly, although early studies on Bernard-Soulier patients reported that platelet aggregation in response to collagen was normal, their transformation into procoagulant platelets was specifically impaired in response to collagen (but not other agonists).(Walsh et al., 1975) More recently, a Bernard-Soulier patient with mutations in both GPIbα and filamin A was also reported to exhibit defects in GPVI-mediated signaling responses.(Li et al., 2015) Although the authors contended that this defect might be due to the filamin A mutation, this may warrant some reappraisal in light of the data presented herein. Like Bernard-Soulier platelets, we found that *GpIbα*^*Δsig/Δsig*^ platelets aggregated normally in response to CRP. The signaling deficit presumably allows sufficient activation of α_IIb_β_3_ for the platelets to aggregate in this assay. This is perhaps unsurprising given that *Gp6*^+/-^ platelet aggregation is only affected at low collagen concentrations.(Kato et al., 2003, Mazharian et al., 2012) Taken together, previous studies support the contention that Bernard-Soulier patient platelets exhibit a partial deficit in GPVI signaling that resembles the deficit in *GpIbα*^*Δsig/Δsig*^ mouse platelets.

The flow assays that we used were highly sensitive to the phenotype of the *GpIbα*^*Δsig/Δsig*^ platelets. However, although these assays are more physiological than many other platelet function assays, their interpretation is complex as these assays are dependent upon multiple parameters, including VWF-GPIbα, collagen-GPVI and shear rate. For these reasons, it is difficult to fully isolate/dissect out the contributions of each signaling pathway/variable to platelet accumulation. Platelets can interact with collagen directly through GPVI and α_2_β_1_, and indirectly via GPIbα binding to VWF, the latter being increasingly important as shear rates rise to first capture the platelets and enable the aforementioned direct interactions to take place.(Nieswandt and Watson, 2003, Ruggeri, 2007) This is demonstrated in wild-type mice, similar to previous reports,(Kuijpers et al., 2004, Kuijpers et al., 2003) by the markedly reduced binding of platelets to collagen in the absence of plasma (and therefore VWF) (Figure 6A-B,D). Residual binding of platelets to collagen with blockade of VWF-GPIbα such as in plasma-free conditions may be attributable partly to direct binding of GPVI with collagen. The importance of VWF-GPIbα binding, and also signaling, appears to be greatest at higher shear rates. Although we demonstrated that *GpIbα*^*Δsig/Δsig*^ platelets bind VWF normally, at the highest shear rate tested (3000s^-1^) we saw the largest effect of the *GpIbα*^*Δsig/Δsig*^ genotype in platelet coverage/accumulation when compared to wild-type mice. VWF-GPIbα binding is particularly important for platelet tethering at high shear. Based on our results, it seems likely that VWF-GPIbα signaling is also important at these shear rates. We contend that VWF-GPIbα platelet priming induces some rapid activation of α_IIb_β_3_, which enable the platelets to better withstand the higher shear rates, prior to their interaction/activation by collagen.

Although most evident at the highest shear rate, *GpIbα*^*Δsig/Δsig*^ platelets exhibited reduced accumulation at all shear rates tested (Figs 5-7, panels C). Given that *GpIbα*^*Δsig/Δsig*^ platelets bind VWF normally, and that the surface coverage on collagen was not significantly altered at 200s^-1^ when compared to wild-type platelets (Figure 7A-B), the deficit in subsequent platelet accumulation must be due to reduced reactivity of *GpIbα*^*Δsig/Δsig*^ platelets. This is supported by the clear importance of α_IIb_β_3_ activation to this assay, demonstrated by the effects of GR144053 on both wild-type and *GpIbα*^*Δsig/Δsig*^ platelets (Figs 6 and 7, panels A-B, D-E). The question remains open as to the precise contribution of VWF-GPIbα versus collagen-GPVI signaling deficits to the phenotype of *GpIbα*^*Δsig/Δsig*^ platelets. Our data suggest that both signaling pathways likely contribute to this.

It is important to note that GPVI-mediated signaling in *GpIbα*^*Δsig/Δsig*^ platelets is diminished, rather than ablated. We endeavored to specifically examine the contribution of GPVI signaling in flow assays by inclusion of the GPVI-blocking JAQ1 antibody. JAQ1 led to a marked decrease in both platelet tethering and accumulation at both 1000s^-1^ and 200s^-1^ shear rates (Figure 6F-I and Figure 7F-I). These assays revealed that there is clear synergy between the roles of VWF-GPIbα and collagen-GPVI in this assay, as disruption of either interaction causes a major reduction in platelet accumulation.

GPVI belongs to the immunoglobulin superfamily and signals via tyrosine kinase phosphorylation pathways. To further investigate the defect in GPVI signaling in *GpIbα*^*Δsig/Δsig*^ platelets, analysis of tyrosine phosphorylation downstream of GPVI revealed that SYK and PLCγ2 phosphorylation was reduced in *GpIbα*^*Δsig/Δsig*^ platelets (Figure 4H-I). This was not due to any alteration in cellular levels of SYK, PLCγ2 (Figure 4I) or GPVI (Figure 2C). Interestingly, the diminished phosphorylation was more pronounced for SYK than for PLCγ2 perhaps highlighting the existence of LAT-independent mechanisms of PLCγ2 phosphorylation.(Pasquet et al., 1999) Based on these findings, we hypothesize that the tail of GPIbα may be important for the docking of signaling molecules such as SYK, LAT and PLCγ2 that are downstream of GPVI on ITAM phosphorylated motif of the FcRγ receptor. The normal binding of PI3K to the intracellular tail of GPIbα may be particularly important in this process. Importantly, activation of *GpIbα*^*Δsig/Δsig*^ platelets via CLEC-2, another receptor that signals via an ITAM motif,(Suzuki-Inoue et al., 2011) was similar to wild-type platelets (Figure S3) suggesting that the function of the GPIbα intracellular tail is more important for GPVI mediated responses.

In summary, we generated a novel GPIbα transgenic mouse in which their platelets bind VWF normally, but the subsequent VWF-GPIbα signaling is disrupted. Intriguingly, these mice clearly reveal the molecular link between GPIbα- and GPVI-mediated signaling in platelets and underscore the cooperative functions of these two major platelet receptors.(Pugh et al., 2010) Various studies have demonstrated that platelets function in diverse system beyond hemostasis. In particular, platelets contribute to the host response to infection and inflammation.(Clark et al., 2007, Deppermann and Kubes, 2016, Jenne and Kubes, 2015, Kapur and Semple, 2016) In these settings, VWF-dependent platelet capture in the absence of vessel injury is recognized as an important step. Our recent work suggests that VWF-GPIbα-dependent platelet priming potentiates the recruitment of neutrophils, which may represent a key early event in the targeting of pathogens, but also in the development of deep vein thrombosis.(Constantinescu-Bercu et al., 2020) The *GpIbα*^*Δsig/Δsig*^ mice now provide an invaluable tool to not only probe the importance of the GPIbα-mediated signaling *in vivo*, in inflammatory diseases such as atherosclerosis and deep vein thrombosis, as well as in the host response to infection, but also to fully decipher the molecular dependency of GPVI signaling upon GPIbα.

## Materials and Methods

### Mice

*GpIbα*^*Δsig/Δsig*^ mice were generated in-house by the Medical Research Council transgenic group at Imperial College using the CRISPR-Cas9 genome editing system (Figure 1). Briefly, pronuclear injections (CBAB6F1) were performed with Cas9 mRNA (75ng/µl), guide RNAs (gRNAs; 25-50ng/µl) and single-strand oligo donor DNA (25-50ng/µl). The donor DNA was generated using the Edit-R HDR donor designer oligo software (Dharmacon) with 50 bp homology arms on the 5’ and 3’ end of the Cas9-DNA nick. PAGE Ultramer DNA Oligos – GGTAAGGCCTAATGGGCGAGTGGGGCCTCTGGTAGCAGGACGGCGACCCTGAGCTCTGAGTCAGGGTCGTGGTCAGGACCTATTGGGCACAGTGGGCATTA-were synthesized by Integrated DNA Technologies (Leuven, Belgium). Embryos were transferred to pseudo-pregnant CBAB6F1 female mice. The two founder mice successfully obtained originated from the same gRNA (CGACCCTGACTCAGAGCTGAGGG) and were bred with C57BL/6 mice. F1 *GpIbα*^*Δ*^*sig/+* mice were bred to obtain *GpIbα*^*Δsig/Δsig*^ mice and *GpIbα*^*+/+*^ littermates were used as controls. Genotyping was performed by PCR amplification of total genomic DNA prepared from ear-punch samples. The *GpIbα* allele (551bp) was detected by PCR using the following forward (FW) and reverse (R) primers: AAGCACTCACACCACAAGCC (FW) and AGTATGAATGAGCGGGAGCC (R) and the sequence confirmed by Sanger sequencing (Genewiz).

### Determination of complete blood counts, platelet counts, surface protein expression and platelet activation

Mice were anaesthetized with ketamine/medetomidine and blood was collected retro-orbitally in 3.8% citrate. Blood was diluted with equal volume of saline and analyzed by the clinical pathology laboratory at Hammersmith Hospital to obtain full blood counts. Platelet counts were determined using precision count beads (Biolegend) and flow cytometry according to the manufacturer’s instructions.

Platelets were washed as previously described with the following modifications.(Salles-Crawley et al., 2014) Blood was diluted in an equal volume of modified Tyrode’s buffer supplemented with prostaglandin E1 (PGE1) and apyrase (both from Sigma) and centrifuged at 150x*g* for 10 mins at room temperature (RT). PRP was subsequently centrifuged at 1000x*g* for 10 mins at RT and three additional centrifugation steps were performed to wash the platelets. Platelets were resuspended at 3×10^5^ platelets/μl in modified Tyrode’s buffer. In experiments using plasma-free blood, red blood cells and leukocytes were separately washed twice in PBS, by centrifugation at 650xg for 10 mins at RT and resuspended in Tyrode’s buffer. Washed platelets were subsequently added to obtain plasma-free blood.

Flow cytometry was performed to analyze the surface expression of GPIbα, GPIbβ, α_IIb_β_3_ and GPVI in platelets from *GpIbα*^*Δsig/Δsig*^ and wild-type littermates using the following antibodies (Abs; Emfret): XiaB2, X488, Leo.H4, and JAQ1, respectively. Whole blood was diluted with modified Tyrode’s buffer (1/20) and stained with Abs for 15 mins at RT before being analysed. Mouse platelets were washed as above and incubated with varying concentrations of agonists - ADP (2-20μM; Labmedics), thrombin (0.02-0.2U/ml; Enzyme Research Laboratories [ERL]), CRP (1-10μg/ml; Cambcol Laboratories), rhodocytin (3 and 300nM; kindly provided by Professor Eble and Dr Hughes) in the presence of 2mM CaCl_2_ for 10 mins at RT. Thereafter, platelets were incubated with JON/A-PE and Wug.E9-FITC Abs for 15 mins at RT to analyze the surface expression of activated α_IIb_β_3_ and P-selectin. Samples were analyzed using a BD LSRFortessa X-20 flow cytometer.

### Platelet aggregometry

Platelet aggregation was assessed by light transmission using the Chronolog 700 aggregometer with continuous stirring at 1,200 rpm at 37°C. Washed platelets were resuspended to a final concentration of 3×10^5^ platelets/μl in modified Tyrode’s buffer and supplemented with 70μg/ml fibrinogen (ERL), 1mM CaCl_2_ and different concentrations of ADP (1-10µM), α-Thrombin (10-50mU/ml) or CRP (0.5-10μg/ml). Platelet aggregation was monitored over 6 mins.

### Platelet spreading

Coverslips were coated with fibrinogen (200μg/ml), CRP (100μg/ml), murine VWF (10μg/ml) or BSA (0.5mg/ml) overnight at 4°C. Coverslips were then blocked with PBS-BSA (5mg/ml) for 1 hour at RT. Washed *GpIbα*^*+/+*^ or *GpIbα*^*Δsig/Δsig*^ mouse platelets were added to the coverslips (150μl/coverslip, 25,000 platelets/μl) in the presence or absence of thrombin (1U/ml) or Botrocetin (2μg/ml) and allowed to adhere for 30 mins – 1 hour, at 37°C. When indicated, platelets were incubated with GR144053 (20µM) for 10 minutes to inhibit α_IIb_β_3_ outside-in signaling prior to stimulation with Botrocetin. Coverslips were then washed with PBS, fixed with 10% formalin, and finally quenched with 50mM NH_4_Cl-PBS. Platelets were then permeabilized in 0.1% Triton-PBS and stained with Flash Phalloidin(tm) Green 488 (2U/ml; Biolegend) for 1.5 hours, at RT. Finally, coverslips were mounted onto slides using ProLong(tm) Gold Antifade Mountant with DAPI (Thermofisher). Spread platelets were visualized using either a Vert.A1 inverted microscope (Zeiss; 40x and 63x air objectives) equipped with ExiBlue camera (Q Imaging) or a confocal microscope (SP5 Leica, 63x objective, z-stack, oil immersion). At least 3 fields of view were analyzed per condition. Surface area of spreading platelets was quantified using Slidebook software 5.0 (3i) and filopodia counted independently by two different researchers.

### Western blotting

For analysis of GPVI and CLEC-2 tyrosine-mediated signaling pathways, washed platelets (3×10^5^ platelets/μl) were stimulated for the indicated time points with 3µg/ml CRP or 30 and 300nM rhodocytin, respectively. Samples were lysed with an equal volume of RIPA buffer (Sigma) supplemented with protease and phosphatase inhibitors (cOmplete, mini and PhosSTOP from Roche). The samples were run under reducing conditions with 4-12% Bolt(tm) Bis-Tris Plus or 4-20% Novex(tm) WedgeWell(tm) Tris-Glycine, 1.0 mm pre-cast gels and proteins were transferred to a nitrocellulose membrane. Membranes were blocked for 1 hour in 3% BSA-TBS and incubated overnight at 4°C with the following primary antibodies: anti-phosphotyrosine 4G10 (Millipore), anti-phosphorylated SYK (pY525/526; Abcam), anti-SYK (D1I5Q; Cell signaling Technology), anti-PLCγ2 (Cell signaling Technology), anti-phosphorylated PLCγ2 (pY1217; Cell signaling Technology), anti-β-actin (Cytoskeleton Inc.), anti-GAPDH (1D4; Novus Biological). The membranes were incubated with horseradish peroxidase-conjugated goat anti-mouse/rabbit secondary Abs (Dako) for 1h at RT and developed using Immobilon(tm) Western Chemiluminescent HRP Substrate (Millipore). Detection and quantification of chemiluminescence intensities were quantified by using Chemidoc™ imaging system. and Image Lab 5.2.1 software (BioRad).

### Flow assays

Murine VWF was expressed in HEK293T cells, purified and quantified as previously described.(De Meyer et al., 2011) VenaFluoro8+ microchannels (Cellix) were coated directly with murine VWF (36.75μg/ml) or collagen (200μg/ml; Labmedics) overnight, at 4°C, in a humidified chamber. Channels were blocked for 1 hour, at RT with HEPES Tyrode’s buffer (134mM NaCl, 0.3mM Na_2_HPO_4_, 2.9mM KCl, 12mM NaHCO_3_, 20mM N-2-hydroxyethylpiperazine-N’-2-ethanesulfonic acid, 5mM Glucose, 1mM MgCl_2,_ pH 7.3) supplemented with 1% BSA.

On the day of the experiments, blood was collected retro-orbitally from *GpIbα*^*Δsig/Δsig*^ mice and wild-type littermates in 100µg/ml Hirudin (Refludan, CSL Behring GmbH) and labelled with Dy-Light 488-conjugated rat anti-mouse GPIbβ (Emfret Analytics, 6µg/ml). When indicated, blood was incubated 5 min prior perfusion with GR144053 (10µM), anti-GPVI JAQ1 or control Rat IgG (Emfret; 20µg/ml). Thereafter, whole blood or plasma-free blood was perfused through the channels at 200-1000s^-1^ using a Mirus pump (Cellix) for 3.5 mins and platelet adhesion/aggregate formation monitored in real-time by fluorescence microscopy (Vert.A1 inverted microscope, Zeiss), using an inverted CCD camera (ExiBlue from Q imaging) operated by the SlideBook™5.0 software. Quantification was performed using SlideBook 5.0 software (3i), to analyze platelet coverage, platelet velocity and thrombus build-up.

### Tail-bleeding assay

Tail bleeding time was performed as described previously.(Salles-Crawley et al., 2014, Crawley et al., 2019) Mice were anaesthetized with ketamine/medetomidine, placed on a heating pad (Harvard Apparatus) at 37°C and a 2 mm segment of the tail was sectioned with a sharp blade. The tail was immediately placed in warm PBS and the time taken for the stream of blood to stop for more than 60 seconds was defined as the bleeding time. To determine the extent of blood loss during the first 10 mins, hemoglobin content was determined by the colorimetric cyanmethemoglobin method using Drabkins reagent and bovine hemoglobin as a standard (Sigma).

### Laser-induced thrombosis model

Thrombus formation was evaluated in the cremaster muscle microcirculation as previously described. (Salles-Crawley et al., 2014, Crawley et al., 2019) Ketamine (75mg/kg) and medetomidine (1mg/kg) was initially given as an intraperitoneally injection. The anesthesia was maintained by giving additional ketamine (12.5mg/kg) every 40 mins. Briefly, Dy-Light 488-conjugated rat anti-mouse GPIbβ Ab (0.15µg/g;Emfret) and Alexa 647-conjugated fibrinogen (5% total fibrinogen; Invitrogen) were administered via a cannula inserted in the jugular vein. Vascular injury was induced by a pulse laser (Ablate!, 3i) focused through a 63X water-immersion objective (65-75% intensity, 5-15 pulses). Thrombus formation was followed in real time for 3 mins after the injury. Median integrated fluorescence intensity over time from platelet or fibrin was determined and analyzed as detailed previously.(Salles-Crawley et al., 2014, Crawley et al., 2019) The operator was blinded to the genotypes during both data acquisition and analysis.

## Acknowledgements

The authors acknowledge the technical assistance of Alisha Miller, Elodie Ndjetehe, Ben Moyon and Zoe Webster from Central Biomedical Services and MRC transgenic group at Imperial College. We thank the LMS/NIHR Imperial Biomedical Research Centre Flow Cytometry Facility for support. We would like to thank Sooriya Soman, Dr Pavarthi Sasikumar, Dr Claire Peghaire at Imperial College for technical assistance, and Nilanthi Karawitage at Imperial College Healthcare NHS trust for the use of the aggregometer. We are grateful to Professor Johannes A. Eble (University of Münster) and Dr Craig E. Hughes (University of Reading) for providing rhodocytin.

This work was supported by the British Heart Foundation grants FS/15/65/32036, PG/17/22/32868 and RG/18/3/33405.

## Authorship Contributions

A.C-B designed and performed experiments, analyzed data and wrote the manuscript; K.J.W. designed and performed experiments and revised the manuscript; P.M and K.V. provided critical reagents and revised the manuscript; J.T.B.C designed experiments, prepared the s and wrote the manuscript; I.I.S-C designed and performed experiments, analyzed data, prepared the s and wrote the manuscript.

## References

Alshehri, O. M., Hughes, C. E., Montague, S., Watson, S. K., Frampton, J., Bender, M. & Watson, S. P. 2015. Fibrin activates GPVI in human and mouse platelets. Blood, 126, 1601–8.

Andrews, R. K. & Berndt, M. C. 2013. Bernard-Soulier syndrome: an update. Semin Thromb Hemost, 39, 656–62.

Arthur, J. F., Gardiner, E. E., Matzaris, M., Taylor, S. G., Wijeyewickrema, L., Ozaki, Y., Kahn, M. L., Andrews, R. K. & Berndt, M. C. 2005. Glycoprotein VI is associated with GPIb-IX-V on the membrane of resting and activated platelets. Thromb Haemost, 93, 716–23.

Baker, J., Griggs, R. K., Falati, S. & Poole, A. W. 2004. GPIb potentiates GPVI-induced responses in human platelets. Platelets, 15, 207–14.

Bergmeier, W., Piffath, C. L., Goerge, T., Cifuni, S. M., Ruggeri, Z. M., Ware, J. & Wagner, D. D. 2006. The role of platelet adhesion receptor GPIbalpha far exceeds that of its main ligand, von Willebrand factor, in arterial thrombosis. Proc Natl Acad Sci U S A, 103, 16900–5.

Clark, S. R., Ma, A. C., Tavener, S. A., Mcdonald, B., Goodarzi, Z., Kelly, M. M., Patel, K. D., Chakrabarti, S., Mcavoy, E., Sinclair, G. D., Keys, E. M., Allen-Vercoe, E., Devinney, R., Doig, C. J., Green, F. H. & Kubes, P. 2007. Platelet TLR4 activates neutrophil extracellular traps to ensnare bacteria in septic blood. Nat Med, 13, 463–9.

Constantinescu-Bercu, A., Grassi, L., Frontini, M., Salles-Crawley, I., Woollard, K. & Crawley, J. T. 2020. Activated alphaIIbbeta3 on platelets mediates flow-dependent NETosis via SLC44A2. Elife, 9.

Crawley, J. T. B., Zalli, A., Monkman, J. H., Petri, A., Lane, D. A., Ahnstrom, J. & Salles-Crawley, I. 2019. Defective fibrin deposition and thrombus stability in Bambi(-/-) mice are mediated by elevated anticoagulant function. J Thromb Haemost, 17, 1935–1949.

Dai, K., Bodnar, R., Berndt, M. C. & Du, X. 2005. A critical role for 14-3-3zeta protein in regulating the VWF binding function of platelet glycoprotein Ib-IX and its therapeutic implications. Blood, 106, 1975–81.

David, T., Strassel, C., Eckly, A., Cazenave, J. P., Gachet, C. & Lanza, F. 2010. The platelet glycoprotein GPIbbeta intracellular domain participates in von Willebrand factor induced-filopodia formation independently of the Ser 166 phosphorylation site. J Thromb Haemost, 8, 1077–87.

De Meyer, S. F., Budde, U., Deckmyn, H. & Vanhoorelbeke, K. 2011. In vivo von Willebrand factor size heterogeneity in spite of the clinical deficiency of ADAMTS-13. J Thromb Haemost, 9, 2506–8.

Denis, C., Methia, N., Frenette, P. S., Rayburn, H., Ullman-Cullere, M., Hynes, R. O. & Wagner, D. D. 1998. A mouse model of severe von Willebrand disease: defects in hemostasis and thrombosis. Proc Natl Acad Sci U S A, 95, 9524–9.

Deppermann, C. & Kubes, P. 2016. Platelets and infection. Semin Immunol, 28, 536–545.

Dubois, C., Panicot-Dubois, L., Gainor, J. F., Furie, B. C. & Furie, B. 2007. Thrombin-initiated platelet activation in vivo is vWF independent during thrombus formation in a laser injury model. J Clin Invest, 117, 953–60.

Dubois, C., Panicot-Dubois, L., Merrill-Skoloff, G., Furie, B. & Furie, B. C. 2006. Glycoprotein VI-dependent and -independent pathways of thrombus formation in vivo. Blood, 107, 3902–6.

Durrant, T. N., Van Den Bosch, M. T. & Hers, I. 2017. Integrin alphaIIbbeta3 outside-in signaling. Blood, 130, 1607–1619.

Estevez, B., Kim, K., Delaney, M. K., Stojanovic-Terpo, A., Shen, B., Ruan, C., Cho, J., Ruggeri, Z. M. & Du, X. 2016. Signaling-mediated cooperativity between glycoprotein Ib-IX and protease-activated receptors in thrombin-induced platelet activation. Blood, 127, 626–36.

Estevez, B., Stojanovic-Terpo, A., Delaney, M. K., O’brien, K. A., Berndt, M. C., Ruan, C. & Du, X. 2013. LIM kinase-1 selectively promotes glycoprotein Ib-IX-mediated TXA2 synthesis, platelet activation, and thrombosis. Blood, 121, 4586–94.

Ezumi, Y., Shindoh, K., Tsuji, M. & Takayama, H. 1998. Physical and functional association of the Src family kinases Fyn and Lyn with the collagen receptor glycoprotein VI-Fc receptor gamma chain complex on human platelets. J Exp Med, 188, 267–76.

Falati, S., Edmead, C. E. & Poole, A. W. 1999. Glycoprotein Ib-V-IX, a receptor for von Willebrand factor, couples physically and functionally to the Fc receptor gamma-chain, Fyn, and Lyn to activate human platelets. Blood, 94, 1648–56.

Feng, S., Resendiz, J. C., Lu, X. & Kroll, M. H. 2003. Filamin A binding to the cytoplasmic tail of glycoprotein Ibalpha regulates von Willebrand factor-induced platelet activation. Blood, 102, 2122–9.

Garcia, A., Quinton, T. M., Dorsam, R. T. & Kunapuli, S. P. 2005. Src family kinase-mediated and Erk-mediated thromboxane A2 generation are essential for VWF/GPIb-induced fibrinogen receptor activation in human platelets. Blood, 106, 3410–4.

Goto, S., Tamura, N., Handa, S., Arai, M., Kodama, K. & Takayama, H. 2002. Involvement of glycoprotein VI in platelet thrombus formation on both collagen and von Willebrand factor surfaces under flow conditions. Circulation, 106, 266–72.

Gu, M., Xi, X., Englund, G. D., Berndt, M. C. & Du, X. 1999. Analysis of the roles of 14-3-3 in the platelet glycoprotein Ib-IX-mediated activation of integrin alpha(IIb)beta(3) using a reconstituted mammalian cell expression model. J Cell Biol, 147, 1085–96.

Jenne, C. N. & Kubes, P. 2015. Platelets in inflammation and infection. Platelets, 26, 286–92.

Ju, L., Chen, Y., Xue, L., Du, X. & Zhu, C. 2016. Cooperative unfolding of distinctive mechanoreceptor domains transduces force into signals. Elife, 5.

Kanaji, T., Russell, S., Cunningham, J., Izuhara, K., Fox, J. E. & Ware, J. 2004. Megakaryocyte proliferation and ploidy regulated by the cytoplasmic tail of glycoprotein Ibalpha. Blood, 104, 3161–8.

Kanaji, T., Russell, S. & Ware, J. 2002. Amelioration of the macrothrombocytopenia associated with the murine Bernard-Soulier syndrome. Blood, 100, 2102–7.

Kapur, R. & Semple, J. W. 2016. Platelets as immune-sensing cells. Blood Adv, 1, 10–14.

Kasirer-Friede, A., Cozzi, M. R., Mazzucato, M., De Marco, L., Ruggeri, Z. M. & Shattil, S. J. 2004. Signaling through GP Ib-IX-V activates alpha IIb beta 3 independently of other receptors. Blood, 103, 3403–11.

Kato, K., Kanaji, T., Russell, S., Kunicki, T. J., Furihata, K., Kanaji, S., Marchese, P., Reininger, A., Ruggeri, Z. M. & Ware, J. 2003. The contribution of glycoprotein VI to stable platelet adhesion and thrombus formation illustrated by targeted gene deletion. Blood, 102, 1701–7.

Kuijpers, M. J., Schulte, V., Bergmeier, W., Lindhout, T., Brakebusch, C., Offermanns, S., Fassler, R., Heemskerk, J. W. & Nieswandt, B. 2003. Complementary roles of glycoprotein VI and alpha2beta1 integrin in collagen-induced thrombus formation in flowing whole blood ex vivo. FASEB J, 17, 685–7.

Kuijpers, M. J., Schulte, V., Oury, C., Lindhout, T., Broers, J., Hoylaerts, M. F., Nieswandt, B. & Heemskerk, J. W. 2004. Facilitating roles of murine platelet glycoprotein Ib and alphaIIbbeta3 in phosphatidylserine exposure during vWF-collagen-induced thrombus formation. J Physiol, 558, 403–15.

Lanza, F. 2006. Bernard-Soulier syndrome (hemorrhagiparous thrombocytic dystrophy). Orphanet J Rare Dis, 1, 46.

Li, J., Dai, K., Wang, Z., Cao, L., Bai, X. & Ruan, C. 2015. Platelet functional alterations in a Bernard-Soulier syndrome patient with filamin A mutation. J Hematol Oncol, 8, 79.

Li, R. 2018. The Glycoprotein Ib-IX-V Complex

Michelson AD, Cattaneo M, Frelinger L, Newman PJ. Elsevier/Academic Press.

Li, Z., Xi, X., Gu, M., Feil, R., Ye, R. D., Eigenthaler, M., Hofmann, F. & Du, X. 2003. A stimulatory role for cGMP-dependent protein kinase in platelet activation. Cell, 112, 77–86.

Li, Z., Zhang, G., Feil, R., Han, J. & Du, X. 2006. Sequential activation of p38 and ERK pathways by cGMP-dependent protein kinase leading to activation of the platelet integrin alphaIIb beta3. Blood, 107, 965–72.

Mangin, P., David, T., Lavaud, V., Cranmer, S. L., Pikovski, I., Jackson, S. P., Berndt, M. C., Cazenave, J. P., Gachet, C. & Lanza, F. 2004. Identification of a novel 14-3-3zeta binding site within the cytoplasmic tail of platelet glycoprotein Ibalpha. Blood, 104, 420–7.

Mangin, P., Yuan, Y., Goncalves, I., Eckly, A., Freund, M., Cazenave, J. P., Gachet, C., Jackson, S. P. & Lanza, F. 2003. Signaling role for phospholipase C gamma 2 in platelet glycoprotein Ib alpha calcium flux and cytoskeletal reorganization. Involvement of a pathway distinct from FcR gamma chain and Fc gamma RIIA. J Biol Chem, 278, 32880–91.

Mangin, P. H., Receveur, N., Wurtz, V., David, T., Gachet, C. & Lanza, F. 2009. Identification of five novel 14-3-3 isoforms interacting with the GPIb-IX complex in platelets. J Thromb Haemost, 7, 1550–5.

Mazharian, A., Wang, Y. J., Mori, J., Bem, D., Finney, B., Heising, S., Gissen, P., White, J. G., Berndt, M. C., Gardiner, E. E., Nieswandt, B., Douglas, M. R., Campbell, R. D., Watson, S. P. & Senis, Y. A. 2012. Mice lacking the ITIM-containing receptor G6b-B exhibit macrothrombocytopenia and aberrant platelet function. Sci Signal, 5, ra78.

Mazzucato, M., Pradella, P., Cozzi, M. R., De Marco, L. & Ruggeri, Z. M. 2002. Sequential cytoplasmic calcium signals in a 2-stage platelet activation process induced by the glycoprotein Ibalpha mechanoreceptor. Blood, 100, 2793–800.

Morowski, M., Vogtle, T., Kraft, P., Kleinschnitz, C., Stoll, G. & Nieswandt, B. 2013. Only severe thrombocytopenia results in bleeding and defective thrombus formation in mice. Blood, 121, 4938–47.

Mu, F. T., Andrews, R. K., Arthur, J. F., Munday, A. D., Cranmer, S. L., Jackson, S. P., Stomski, F. C., Lopez, A. F. & Berndt, M. C. 2008. A functional 14-3-3zeta-independent association of PI3-kinase with glycoprotein Ib alpha, the major ligand-binding subunit of the platelet glycoprotein Ib-IX-V complex. Blood, 111, 4580–7.

Nakamura, F., Pudas, R., Heikkinen, O., Permi, P., Kilpelainen, I., Munday, A. D., Hartwig, J. H., Stossel, T. P. & Ylanne, J. 2006. The structure of the GPIb-filamin A complex. Blood, 107, 1925–32.

Nesbitt, W. S., Kulkarni, S., Giuliano, S., Goncalves, I., Dopheide, S. M., Yap, C. L., Harper, I. S., Salem, H. H. & Jackson, S. P. 2002. Distinct glycoprotein Ib/V/IX and integrin alpha IIbbeta 3-dependent calcium signals cooperatively regulate platelet adhesion under flow. J Biol Chem, 277, 2965–72.

Nieswandt, B., Brakebusch, C., Bergmeier, W., Schulte, V., Bouvard, D., Mokhtari-Nejad, R., Lindhout, T., Heemskerk, J. W., Zirngibl, H. & Fassler, R. 2001. Glycoprotein VI but not alpha2beta1 integrin is essential for platelet interaction with collagen. EMBO J, 20, 2120–30.

Nieswandt, B., Pleines, I. & Bender, M. 2011. Platelet adhesion and activation mechanisms in arterial thrombosis and ischaemic stroke. J Thromb Haemost, 9 Suppl 1, 92–104.

Nieswandt, B. & Watson, S. P. 2003. Platelet-collagen interaction: is GPVI the central receptor? Blood, 102, 449–61.

Pasquet, J. M., Gross, B., Quek, L., Asazuma, N., Zhang, W., Sommers, C. L., Schweighoffer, E., Tybulewicz, V., Judd, B., Lee, J. R., Koretzky, G., Love, P. E., Samelson, L. E. & Watson, S. P. 1999. LAT is required for tyrosine phosphorylation of phospholipase cgamma2 and platelet activation by the collagen receptor GPVI. Mol Cell Biol, 19, 8326–34.

Pugh, N., Simpson, A. M., Smethurst, P. A., De Groot, P. G., Raynal, N. & Farndale, R. W. 2010. Synergism between platelet collagen receptors defined using receptor-specific collagen-mimetic peptide substrata in flowing blood. Blood, 115, 5069–79.

Rayes, J., Watson, S. P. & Nieswandt, B. 2019. Functional significance of the platelet immune receptors GPVI and CLEC-2. J Clin Invest, 129, 12–23.

Ruggeri, Z. M. 2007. The role of von Willebrand factor in thrombus formation. Thromb Res, 120 Suppl 1, S5–9.

Salles-Crawley, I., Monkman, J. H., Ahnstrom, J., Lane, D. A. & Crawley, J. T. 2014. Vessel wall BAMBI contributes to hemostasis and thrombus stability. Blood, 123, 2873–81.

Schmaier, A. A., Zou, Z., Kazlauskas, A., Emert-Sedlak, L., Fong, K. P., Neeves, K. B., Maloney, S. F., Diamond, S. L., Kunapuli, S. P., Ware, J., Brass, L. F., Smithgall, T. E., Saksela, K. & Kahn, M. L. 2009. Molecular priming of Lyn by GPVI enables an immune receptor to adopt a hemostatic role. Proc Natl Acad Sci U S A, 106, 21167–72.

Schoenwaelder, S. M., Darbousset, R., Cranmer, S. L., Ramshaw, H. S., Orive, S. L., Sturgeon, S., Yuan, Y., Yao, Y., Krycer, J. R., Woodcock, J., Maclean, J., Pitson, S., Zheng, Z., Henstridge, D. C., Van Der Wal, D., Gardiner, E. E., Berndt, M. C., Andrews, R. K., James, D. E., Lopez, A. F. & Jackson, S. P. 2016. 14-3-3zeta regulates the mitochondrial respiratory reserve linked to platelet phosphatidylserine exposure and procoagulant function. Nat Commun, 7, 12862.

Stalker, T. J. 2020. Mouse laser injury models: variations on a theme. Platelets, 31, 423–431.

Suzuki-Inoue, K., Inoue, O. & Ozaki, Y. 2011. Novel platelet activation receptor CLEC-2: from discovery to prospects. J Thromb Haemost, 9 Suppl 1, 44–55.

Tsuji, M., Ezumi, Y., Arai, M. & Takayama, H. 1997. A novel association of Fc receptor gamma-chain with glycoprotein VI and their co-expression as a collagen receptor in human platelets. J Biol Chem, 272, 23528–31.

Walsh, P. N., Mills, D. C., Pareti, F. I., Stewart, G. J., Macfarlane, D. E., Johnson, M. M. & Egan, J. J. 1975. Hereditary giant platelet syndrome. Absence of collagen-induced coagulant activity and deficiency of factor-XI binding to platelets. Br J Haematol, 29, 639–55.

Ware, J., Russell, S. & Ruggeri, Z. M. 2000. Generation and rescue of a murine model of platelet dysfunction: the Bernard-Soulier syndrome. Proceedings of the National Academy of Sciences of the United States of America, 97, 2803–8.

Wu, Y., Suzuki-Inoue, K., Satoh, K., Asazuma, N., Yatomi, Y., Berndt, M. C. & Ozaki, Y. 2001. Role of Fc receptor gamma-chain in platelet glycoprotein Ib-mediated signaling. Blood, 97, 3836–45.

Yin, H., Liu, J., Li, Z., Berndt, M. C., Lowell, C. A. & Du, X. 2008. Src family tyrosine kinase Lyn mediates VWF/GPIb-IX-induced platelet activation via the cGMP signaling pathway. Blood, 112, 1139–46.

Yin, H., Stojanovic-Terpo, A., Xu, W., Corken, A., Zakharov, A., Qian, F., Pavlovic, S., Krbanjevic, A., Lyubimov, A. V., Wang, Z. J., Ware, J. & Du, X. 2013. Role for platelet glycoprotein Ib-IX and effects of its inhibition in endotoxemia-induced thrombosis, thrombocytopenia, and mortality. Arterioscler Thromb Vasc Biol, 33, 2529–37.

Zhang, W., Deng, W., Zhou, L., Xu, Y., Yang, W., Liang, X., Wang, Y., Kulman, J. D., Zhang, X. F. & Li, R. 2015. Identification of a juxtamembrane mechanosensitive domain in the platelet mechanosensor glycoprotein Ib-IX complex. Blood, 125, 562–9.

