## Supplementary Figures and table for "The GPIbα intracellular tail - role in transducing VWF- and Collagen/GPVI-mediated signaling"

**SUPPLEMENTARY INFORMATION**

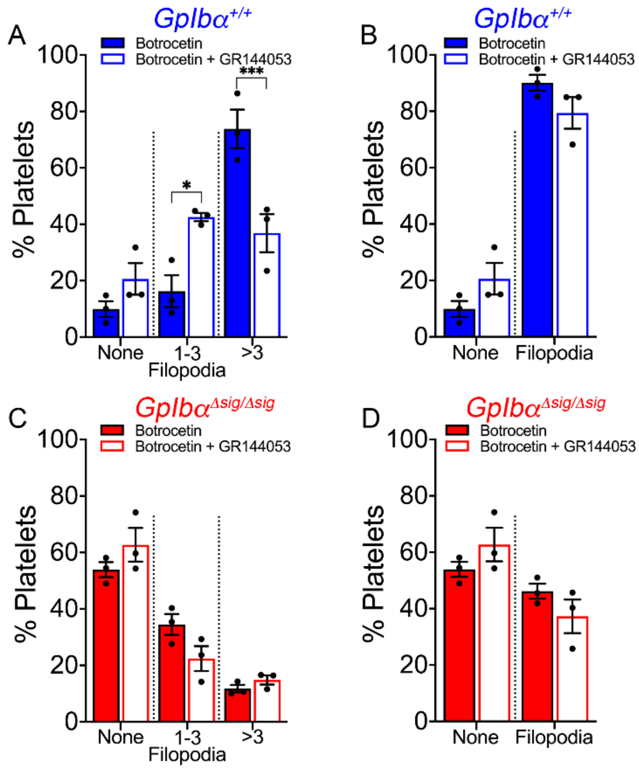


**Figure S1: *GpIbα*$^{Δsig/Δsig}$ platelets exhibit disrupted GPIbα-mediated signaling.** *GpIbα*$^{+/+}$ (A,B, blue bars) and *GpIbα*$^{Δsig/Δsig}$ (C,D, red bars) platelets (n=3 for each genotype with individual data points representing the average of 3 fields of view) were spread on murine VWF and stained with Phalloidin-Alexa 488, in the presence of Botrocetin supplemented or not with GR144053. (A,C) Percentage of platelets with no filopodia, 1-3 filopodia or >3 filopodia formed on murine VWF upon stimulation with Botrocetin or Botrocetin and GR144053. (B,D) Percentage of platelets with or without filopodia formed on murine VWF upon stimulation with Botrocetin or Botrocetin and GR144053. All data is shown as mean ± SEM and was analyzed using two-way ANOVA followed by Sidak's multiple comparison test; *p<0.05, ***p<0.001. Also see Figure 2G-J.

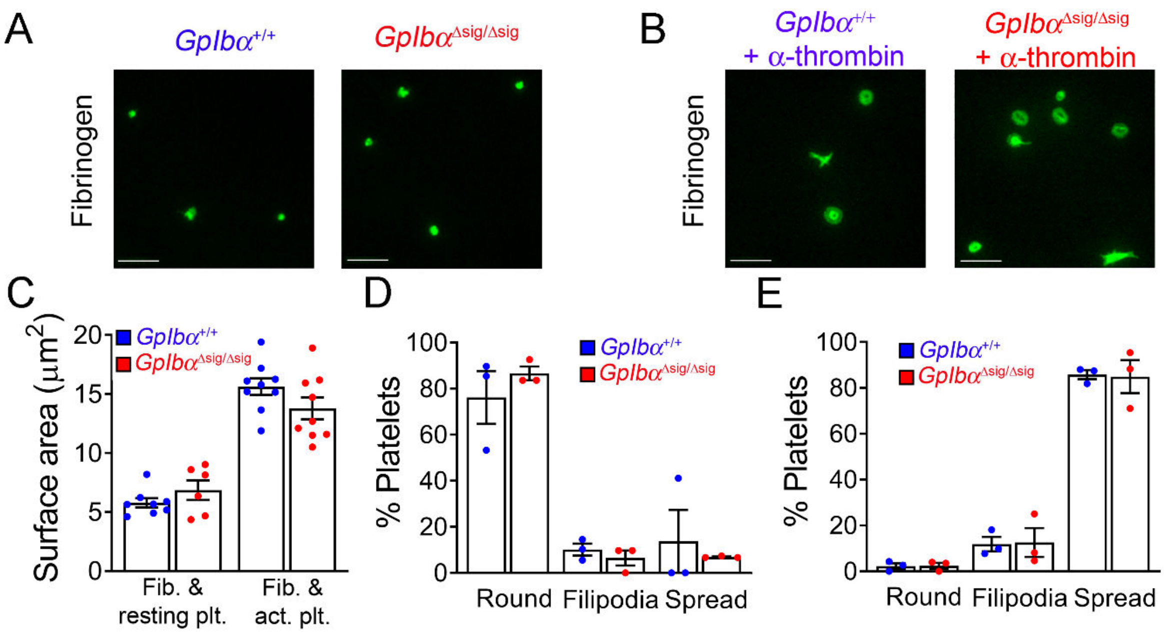


**Figure S2: $GpIb\alpha^{\Delta sig/\Delta sig}$ platelets spread normally on fibrinogen under basal and stimulated conditions.** Representative micrographs (n=3 for each genotype; 3 fields of view analyzed per condition; scale bar 10 µm) of $GpIb\alpha^{+/+}$ and $GpIb\alpha^{\Delta sig/\Delta sig}$ platelets in the absence (A) or presence of 0.2U/ml α-thrombin (B) and spread on fibrinogen. Platelet spreading was visualized by Phalloidin-Alexa 488 staining. Bar graphs quantifying the surface area (C) and percentages of platelets that remained round, formed filopodia or spread on fibrinogen under basal conditions (D) or activated with α-thrombin (E). The data represent the mean ± SEM and was analyzed using two-way ANOVA followed by Sidak's multiple comparison test; $p>0.05$. Fib.:fibrinogen; plt.: platelets; act.: α-thrombin-activated. Also see Figure 4A-G.

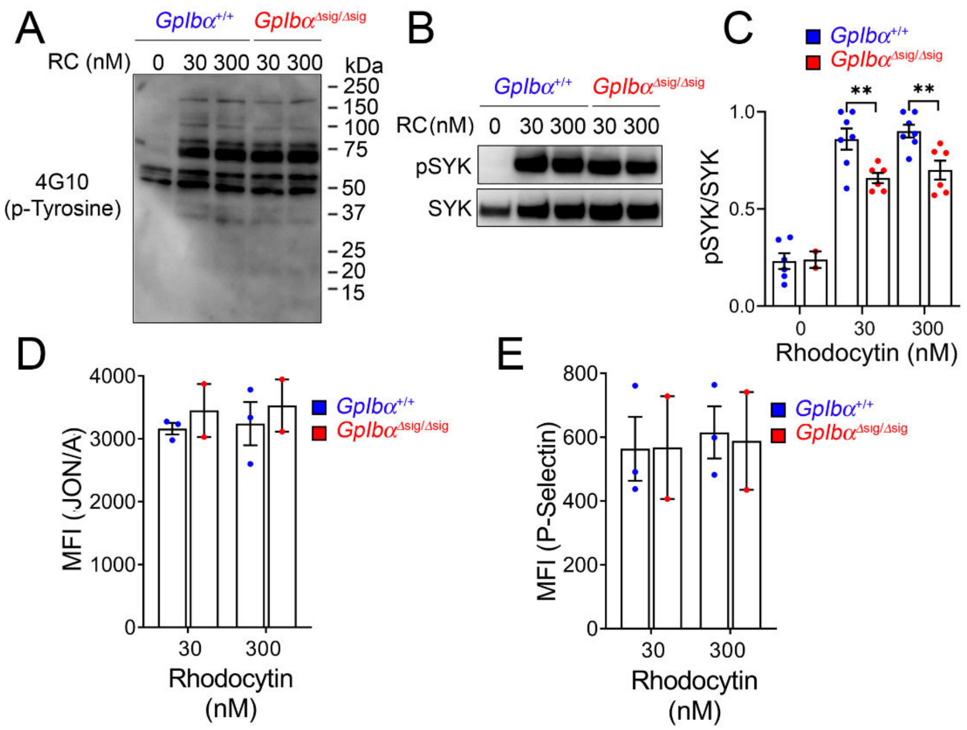


**Figure S3: Truncation of the GPIbα intracellular tail does not greatly influence CLEC-2 mediated signaling.** (A) Western blot analyzing tyrosine kinase phosphorylation in platelets from *GpIbα*$^{+/+}$ and *GpIbα*$^{\Delta sig/\Delta sig}$ mice, following 5 min stimulation with rhodocytin (RC; 30 and 300nM), (representative of n=3). (B) Western blot and (C) bar graph analyzing the levels of phosphorylated and non-phosphorylated SYK in platelets from *GpIbα*$^{+/+}$ and *GpIbα*$^{\Delta sig/\Delta sig}$ mice, after 3 mins stimulation with RC (representative of n=3). (D-E) Flow cytometric analysis of surface expression of activated $\alpha_{IIb}\beta_3$ (D) and P-selectin (E) in *GpIbα*$^{+/+}$ and *GpIbα*$^{\Delta sig/\Delta sig}$ platelets (n=2-3) after stimulation with rhodocytin (RC, 30-300nM). Data is shown as mean ± SEM and analyzed using two-way ANOVA followed by Sidak's multiple comparison test; **p<0.001. Also see Figure 4H-K.

**Table S1** Hematological parameters

| | $GpIb\alpha^{+/+}$ | $GpIb\alpha^{\Delta sig/\Delta sig}$ |
|---|---|---|
| PLT ($10^3$/µl) | 1028 ± 187 | 818 ± 188**** |
| RBC ($10^6$/µl) | 9.0 ± 1.0 | 8.8 ± 0.7 |
| HCT (%) | 50.7 ± 5.2 | 49.7 ± 3.2 |
| WBC ($10^3$/µl) | 5.9 ± 1.8 | 6.7 ± 1.5 |

PLT, platelets; RBC, red blood cells; HCT, hematocrit, WBC, white blood cells; ****P <0.001, unpaired, two-tailed t-test, mean ± SD (n=10 per genotype)

**Video 1 (separate file). Laser-induced thrombus formation in a $GpIb\alpha^{+/+}$ and $GpIb\alpha^{\Delta sig/\Delta sig}$ mouse:** Representative videos of fluorescently-labeled platelets (green) and fibrin(ogen) (red) accumulating at the site of laser-induced injury in a cremaster muscle arteriole of a $GpIb\alpha^{+/+}$ and $GpIb\alpha^{\Delta sig/\Delta sig}$ mouse. Thrombus formation was studied using a combination of brightfield and fluorescence microscopy. Results are presented in Figure 3. A timer is shown in the top left corner (hh:mm:ss:000) and a 10 µm scale bar in the bottom left corner.

**Video 2 (separate file). Platelet capture on murine VWF-coated microchannels.** Representative videos of hirudin-anticoagulated blood from a $GpIb\alpha^{+/+}$ and $GpIb\alpha^{\Delta sig/\Delta sig}$ mouse perfused over mouse VWF. Thrombus formation was visualized over 3 minutes of perfusion at 1000s-$^{1}$. Results are presented in Figure 2D-F. A timer is shown in the top left corner (hh:mm:ss:000) and a 10 µm scale bar in the bottom left corner.

**Video 3 (separate file). Formation of platelet aggregates on collagen-coated microchannels at 3000s$^{-1}$.** Representative video of hirudin-anticoagulated blood from a $GpIb\alpha^{+/+}$ and $GpIb\alpha^{\Delta sig/\Delta sig}$ mouse over fibrillar collagen type I. Thrombus formation was visualized over 3 minutes of perfusion at 3000s-$^{1}$. Results are presented in Figure 5. A timer is shown in the top left corner (hh:mm:ss:000) and a 10 µm scale bar in the bottom left corner.

**Video 4 (separate file). Formation of platelet aggregates on collagen-coated microchannels at 1000s$^{-1}$.** Representative video of hirudin-anticoagulated blood from a $GpIb\alpha^{+/+}$ and $GpIb\alpha^{\Delta sig/\Delta sig}$ mouse over fibrillar collagen type I. Thrombus formation was visualized over 3 minutes of perfusion at 1000s-$^{1}$. Results are presented in Figure 6. A timer is shown in the top left corner (hh:mm:ss:000) and a 10 µm scale bar in the bottom left corner.

**Video 5 (separate file). Formation of platelet aggregates on collagen-coated microchannels at 200s$^{-1}$.** Representative video of hirudin-anticoagulated blood from a $GpIb\alpha^{+/+}$ and $GpIb\alpha^{\Delta sig/\Delta sig}$ mouse over fibrillar collagen type I. Thrombus formation was visualized over 3 minutes of perfusion at 200s-$^{1}$. Results are presented in Figure 7. A timer is shown in the top left corner (hh:mm:ss:000) and a 10 µm scale bar in the bottom left corner.
